# Maturation-dependent structural dynamics and nanomechanical properties of dengue virus revealed by high-speed AFM and 3D force mapping

**DOI:** 10.64898/2026.05.07.723438

**Authors:** Steven J. McArthur, Kenichi Umeda, Noriyuki Kodera

## Abstract

Although the maturation state of dengue virus (DENV) particles is a key determinant of their infectivity, maturation is unusually inefficient. Fully mature and immature DENV particles are well-studied; however, little is known about partially mature particles. Moreover, single-particle structural dynamics and nanomechanical properties are unknown. Here, we observe wildtype and immature DENV particles using a single-particle approach combining high-speed AFM (HS-AFM) and 3D force mapping (3DFM). HS-AFM shows that the conformations of each morphotype are heterogeneous and dynamic in liquid, particularly partially mature virions. Tracking immature prM–E spikes elucidates their dynamic movements, which show intraviral variation and constrained independence. 3DFM measurements suggest that internal DENV structure is also heterogeneous and undergoes maturation-dependent changes, with the nucleocapsid core not occupying the full internal volume of immature virions. This approach complements current structural virology techniques and adds a new dimension to our understanding of the structural properties of viruses.

## MAIN

One of the most challenging aspects of developing therapeutic and prophylactic strategies targeting dengue virus (DENV) has been its remarkable structural plasticity and dynamic nature^1–5^. The DENV virion is a particle of approximately 50 nm, composed of a capsid-bound RNA genome (nucleocapsid) enveloped in a phospholipid bilayer in which the glycoproteins prM and E are embedded. The prM–E heterocomplex has been shown to adopt a wide range of conformations, depending on pH, temperature, and maturation state^6–8^. Maturation refers to the proteolytic cleavage of prM by the host enzyme furin to yield the soluble fragment pr and the transmembrane fragment M; this event is nominally required for infection^9,10^. Strangely, DENV maturation is inefficient, even compared to other *Flaviviridae*^11^. This inefficiency leads to a spectrum of heterogeneous morphotypes, since immature (IM) prM–E forms tripods or spikes that protrude from the virion surface whereas mature (MA) M–E forms a smooth herringbone-like pattern along the virion surface (Fig. 1a)^12–15^. Notably, cryo-electron microscopy (cryoEM) studies have shown that partially mature (PM) DENV present both spiky and smooth forms on a single virion, forming ‘mosaic’ particles^16,17^.

**Figure 1.**
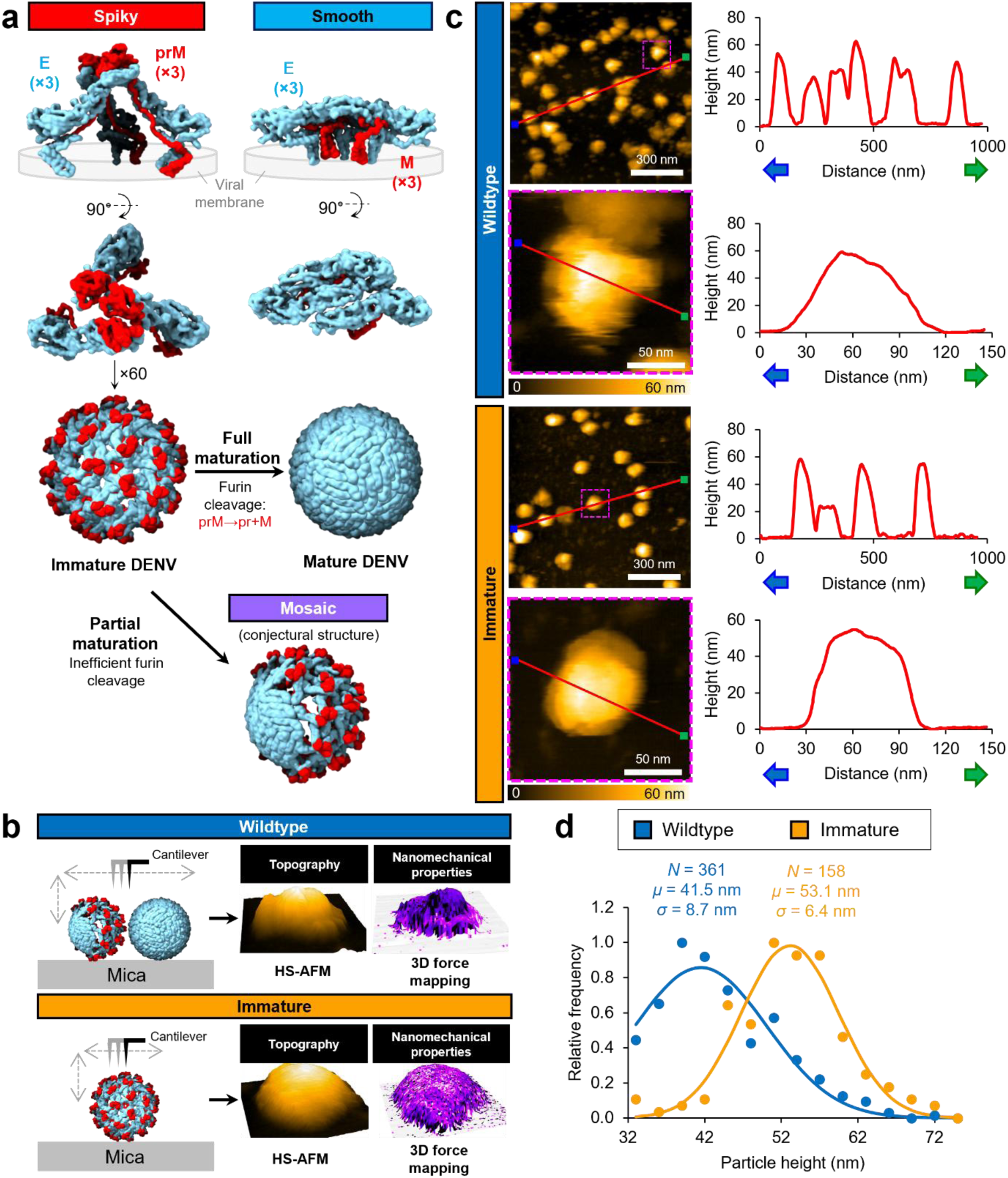
Preparation and characterization of wildtype and immature DENV samples. (**a**) Immature prM–E complexes associate as trimeric spikes in the viral membrane (PDB 4B03). Mature M–E are found in a smooth, herringbone-like conformation lying along the surface of the viral membrane (PDB 4CCT). 60 prM–E spikes associate to form an icosahedrally symmetrical immature virion. Maturation, whereby the host protease furin cleaves prM to produce pr and M fragments, evinces a spiky to smooth conformational change. Furin-mediated cleavage is inefficient, leading to many partially mature virions in which spiky and smooth units are both present (conjectural structure). (**b**) Two sample types analyzed by HS-AFM and 3D force mapping (3DFM): wildtype, containing mature and partially mature virions, and immature, containing immature virions. Red: prM/M; blue: E. (**c**) Representative scans of virus-sized particles from wildtype and immature culture respectively, showing a 1000 × 1000 nm^2^ overview scan (200 × 200 pixels, scan speed 40 µm/s, frame time 10.3 s in trace-retrace mode) and a single virus-sized particle in a 150 × 150 nm^2^ scan (150 × 150 pixels, scan speed 20 µm/s, frame time 1.5 s in only-trace mode). Line profiles along the red lines from blue end to green end are shown at right. (**d**) Frequency distribution of particle heights greater than 33 nm.

While static structures of IM and MA virions have been resolved, our understanding of DENV conformational diversity and dynamics has been limited^1^. Time-resolved Förster resonance energy transfer and hydrogen–deuterium exchange mass spectrometry have both been applied, robustly demonstrating that population-averaged DENV are structurally dynamic in liquid^18–20^; however, the dynamics of individual virions remain enigmatic, and the comparative dynamics of different maturation states have not been studied. These dynamics have important biological implications since the exposure of cryptic epitopes has been linked to the formation of cross-reactive non-neutralizing antibodies, which can promote infection by antibody-dependent enhancement (ADE)^21–23^. Moreover, structures of PM virions remain elusive despite comprising up to 42% of DENV progeny *in vitro*^24^ since their heterogeneous nature renders them unsuitable for the ensemble averaging techniques used in structural virology^25^.

This technical limitation has also precluded analysis of the disordered nucleocapsid core. Young’s modulus (YM) and stiffness are viral nanomechanical properties with links to viral adaptability and fitness^26,27^. The biological importance of nanomechanical properties has been demonstrated for other viruses^28,29^: for example, the stiffness of HIV-1 is maturation-dependent^30–33^. We believe these parameters present an unexplored avenue to probe DENV nucleocapsid structure. Thus, to begin to address these unknowns, we present the application of high-speed AFM (HS-AFM) and 3D force mapping (3DFM) (Fig. 1b) in a single-particle approach to characterize the real-time structural dynamics and nanomechanical properties of individual DENV virions in different maturation states.

### Topographical characterization of maturation morphotypes

We cultured DENV serotype 2 (DENV-2), responsible for the highest pooled mortality rate among DENV serotypes^34^, in Vero cells. We isolated two populations of virions: wildtype (WT), and IM samples isolated from cells cultured in medium supplemented with ammonium chloride (Supplementary Fig. 1a), which blocks maturation (Supplementary Fig. 1b–c)^7,35^. Wide-area HS-AFM scans of WT fractions showed an abundance of virus-sized (∼50 nm) spheroid-like particles in the fraction that had the highest infectivity before inactivation; this population was reduced in fractions with lower infectivity (Supplementary Fig. 2), suggesting these particles were virions. Both WT and IM preparations yielded many particles with virus-like heights and widths (Fig. 1c). Overall, WT virus-sized particles showed an average particle height of 41.5 ± 8.7 nm (all values mean ± SD, *n* = 361), whereas IM samples showed a reduction in smaller particles and an increase in larger particles, with an average particle height of 53.1 ± 6.4 nm (*n* = 158) (Fig. 1d). The increase in average particle height in IM samples is consistent with the expected increase in abundance of the larger-diameter spiky morphotype and reduction in the smaller-diameter smooth morphotype^36^, further supporting the identification of these particles as virions. Interestingly, the appearance of individual virions in WT and IM samples was heterogeneous: obviously damaged or defective particles notwithstanding, some degree of variability or imperfection was found in most particles.

To form a basis for morphotype classification, we performed simulated AFM^37^ on the published cryoEM structures of MA and IM virions, as well as an arbitrary PM hybrid structure based on the current understanding of mosaic-type virions (Fig. 1a)^16,38^. We found that the smooth, mosaic, and spiky morphotypes are expected to be observable by AFM (Fig. 2a left). Single-particle HS-AFM scans of virions from WT and IM samples revealed morphologies that corresponded with the simulations (Fig. 2A right). IM samples showed spiky virions, while WT samples showed a mixture of MA-like smooth and PM-like mosaic virions. Although the apparent width of IM virions was increased more than predicted by the simulation, the peak-to-trough prominence of single prM–E spikes (2.6 nm) and base diameter (14.1 nm) corresponded well with the simulation (2.4 and 13.8 nm respectively; Extended Data Fig. 1)^39^. We confirmed that the average maximum height of particles varied significantly according to morphotype, with smooth virions at 41.1 ± 6.6 nm (*n* = 11), mosaic virions at 51.4 ± 6.9 nm (*n* = 35), and spiky virions at 58.5 ± 6.9 nm (*n* = 40) (Fig. 2b).

**Figure 2.**
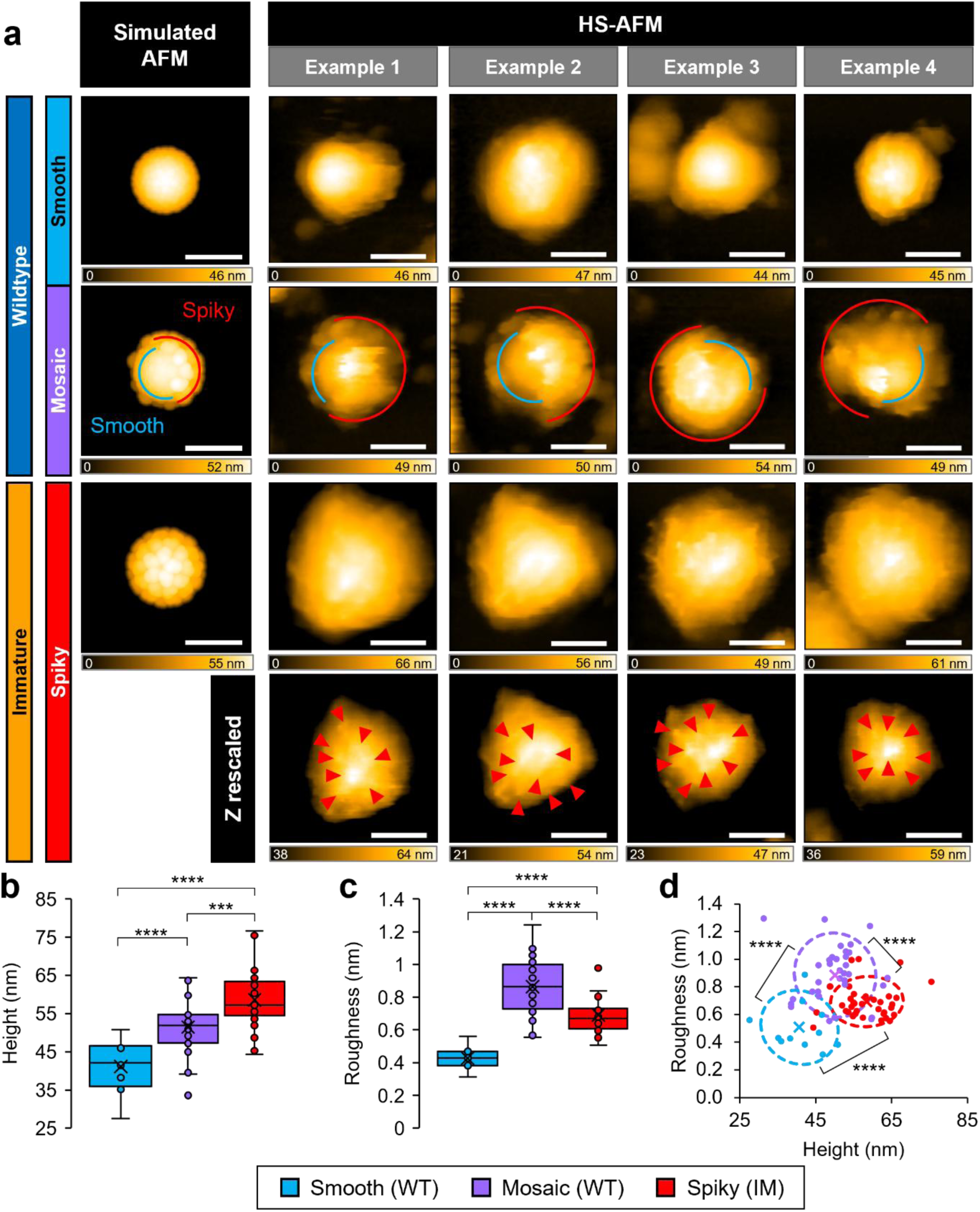
The appearance and topographical properties of DENV virions vary with their maturation state. (**a**) Simulated AFM using the structures from Fig. 1a (left column); and representative experimental HS-AFM images (right four columns), showing particles with smooth-like, mosaic-like, and spiky-like morphologies (scan area 150 × 150 nm^2^ at 150 × 150 pixels, scan speed 20 µm/s, frame time 1.5 s in only-trace mode). Red and blue arcs shown on mosaic virions highlight spiky-like (red) and smooth-like (blue) regions. Bottom row: the four immature virions with height (Z) coloring rescaled to enhance contrast on the virion surface, clarifying spike-like protrusions (red arrowheads). Image processing: tilt-corrected 3-frame triangular average with Gaussian filter (*σ* = 1.0), median effect filter 1 × 1 pixels. Scale bars: 50 nm. (**b**–**c**) Average maximum height (**b**) and surface roughness (**c**) of IM and WT virions. IM: *n* = 40 over 1773 total frames; WT: *n* = 46 over 1621 total frames. Boxes: interquartile range; lines: median; crosses, mean; error bars: SD. ***, *p* < 0.001; ****, *p* < 0.0001 by Welch’s t-test. (**d**) 2D plot of roughness vs. height; ****, *p* < 0.0001 by Hotelling’s t^2^-test. Crosses, mean; dashed ellipses: 2D SD. (**b**–**d**) Each point represents one virion observed over time in a single HS-AFM scan. Orange: IM; blue: WT.

In order to quantitatively compare the topography of the three morphotypes, we next calculated their surface roughness. We confirmed that spiky virions showed increased roughness compared to smooth virions (0.69 ± 0.12 nm and 0.42 ± 0.08 nm respectively) (Fig. 2c). Mosaic virions were rougher than both (0.86 ± 0.18 nm) due to the contrast between spiky and smooth regions as well as instability at the spiky–smooth interface (Supplementary Fig. 3). Considering both height and roughness together, virions were found distributed among three clusters, consisting of low-roughness low-height particles (smooth), high-roughness variable-height particles (mosaic), and medium-roughness large-height particles (spiky) (Fig. 2d).

### Real-time structural dynamics and spike tracking

Using continuous single-virion HS-AFM scanning, we sought to characterize the structural dynamics of each DENV morphotype. Qualitatively, we found that smooth WT virions were stable over time, with occasional small-scale shifts (Fig. 3a top, Supplementary Video 1). Similarly, spiky IM virions tended to be stable overall with occasional small movements (Fig. 3a bottom, Supplementary Video 2). In contrast, mosaic WT particles showed more variation. Many exhibited highly flexible regions that appeared as a smear during HS-AFM imaging (Fig. 3a middle, Supplementary Videos 3–4). Some also showed frequent small rapid movements near the spiky–smooth interface, such as single spikes moving between flat smooth-like and raised spiky-like orientations (Fig. 3a middle, Supplementary Video 3). In contrast, some mosaic WT virions appeared to be more stable, showing only small-scale shifts similar to those seen in MA and IM virions (Supplementary Video 5).

**Figure 3.**
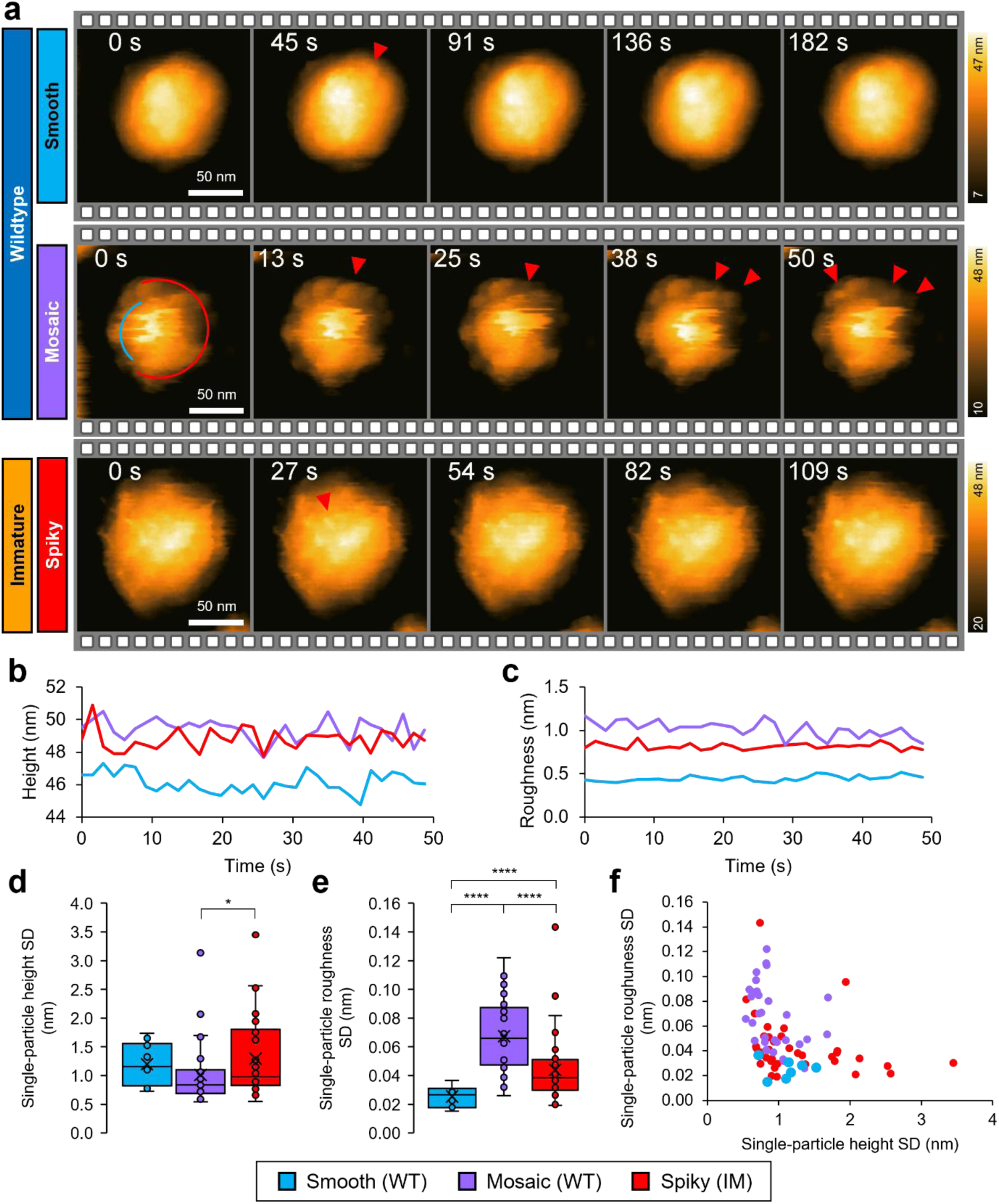
Single-particle HS-AFM reveals maturation-dependent structural dynamics of DENV-2. (**a**) Representative single particles corresponding to each morphotype were observed by HS-AFM (scan area 150 × 150 nm^2^ at 150 × 150 pixels, scan speed 20 µm/s, frame time 1.5 s in only-trace mode). Notable protein movements are indicated by red arrowheads. Scale bars: 50 nm. Image processing: tilt-corrected with Gaussian filter *σ* = 1.0, median effect filter *r* = 1 pixel. (**b**–**c**) Representative particle height (**b**) and roughness (**c**) traces for the single particles shown in panel A over 50 s of continuous HS-AFM observation. See Supplementary Videos 1–3 for the full HS-AFM scans represented in panel A. (**d**–**e**) Summary of standard deviations in the maximum height (**d**) and surface roughness (**e**) of single particles. (**f**) 2D plot of roughness SD vs. height SD for single particles. (**d**–**f**) Each point represents the SD of one height or roughness trace for one particle over time in one continuous HS-AFM scan. Smooth, *n* = 7; mosaic, *n* = 31; spiky, *n* = 36; *, *p* < 0.05; ****, *p* < 0.0001 by Welch’s t-test.

We found no trend in particle height or roughness over time, indicating that virions were stable during HS-AFM imaging (Fig. 3b–c, Extended Data Fig. 2a). Single-particle standard deviations in height over time was also observed to be roughly similar across morphotypes (Fig. 3d). Interestingly, single-particle standard deviations in roughness over time showed morphotype-specific differences, with mosaic virions exhibiting greater roughness variation (0.067 ± 0.026 nm) compared to spiky (0.044 ± 0.024 nm) or smooth (0.026 ± 0.008 nm) (Fig. 3e). Unlike height, which evaluates a single pixel (i.e. the maximum) per particle per frame, roughness compares the topography over all pixels in a region, providing a quantitative assessment of small-scale conformational shifts. Thus, mosaic virions show a higher degree of topographical variation over time than spiky virions, which in turn are more variable than smooth virions. Considering height and roughness SD over time together, spiky particles skew towards higher height variation while mosaic particles skew towards greater roughness variation, illustrating the differences in dynamics among morphotypes (Fig. 3f).

To obtain further insight into the dynamics of DENV, we performed sub-particle analysis of spiky IM virions, whose prM–E spikes can be tracked during continuous HS-AFM observation (Fig. 4a–c, Extended Data Figs. 3–4). Comparison of flattened HS-AFM images with flattened simulated AFM of cryoEM structures^40–42^ showed that the geometric arrangement of prM–E spikes does not precisely conform, with large variations in the gap size between nearby spikes. On average, HS-AFM spike gaps were 6.8% larger (5.5 ± 1.4 nm) than in the cryoEM structure (Fig. 4d). Tracking spikes over time (Fig. 4e, Supplementary Video 6) revealed that framewise displacement from the average position of each spike was similar (1.6 ± 0.4 nm) (Fig. 4f). However, intraviral variation in frame-to-frame velocity (Fig. 4g) and radius of confinement (Fig. 4h) was observed, suggesting some independence of movement within the constraints of the icosahedral lattice. Spearman correlation analysis for frame-to-frame velocity and movement angle among pairs of spikes supported this independence, with only 4 and 1 out of 45 spike pairs showing a moderate positive correlation in velocity and angle respectively (Fig. 4i). Interestingly, comparison of the framewise movement angle distributions for each spike showed that each has a different preferred axis of movement (Fig. 4j). Similar analysis on the spikes of two additional IM virions showed similar results (Extended Data Figs. 3–4). Overall, this suggests that, in a biological-like setting, each spike has its own distinct dynamic motion within the bounds of the lattice; this contrasts with the static symmetrical capsids seen in many non-enveloped viruses^28,29,43^.

**Figure 4.**
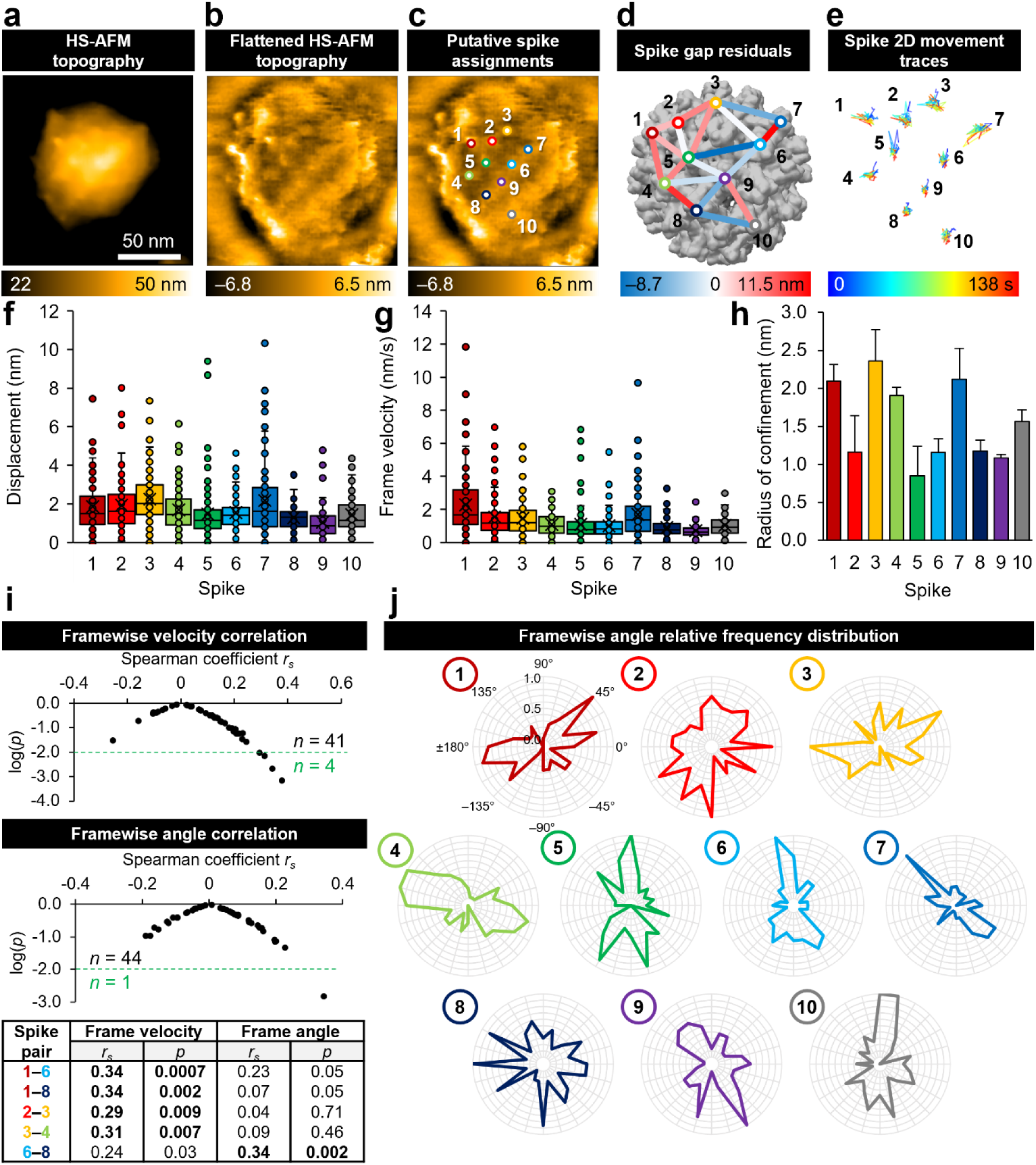
Spikes undergo heterogeneous dynamic movements on the surface of IM-DENV-2. (**a**) Representative HS-AFM image (acquired and processed as described in Fig. 3; scale bar 50 nm). Full HS-AFM scan shown in Supplementary Video 2. (**b**) The same image, processed with a Gaussian filter (σ = 1.0), second-order flattening filters (rows and columns respectively), and a first-order flattening filter in the XY plane. (**c**) The same image, with trackable spikes labeled and arbitrarily numbered. (**d**) Trackable spikes mapped onto the cryoEM structure (PDB 4B03). Residuals in the 3D distance between adjacent spikes in the cryoEM structure and HS-AFM structure are indicated. (**e**) Spike localization traces, coloured blue to red over time. Full traces shown along with flattened HS-AFM scan in Supplementary Video 6. (**f**) Displacement of each spike at each frame relative to the average position of the spike (unresolved frames excluded). (**g**) Spike velocity measured between successive frames (unresolved frames excluded). (**f**–**g**) Boxes: interquartile range; lines: median; crosses, mean; error bars: SD. (**h**) 2D apparent radius of confinement for each spike, calculated using the mean square displacement method. Error bars: SD. (**i**) Pairwise Spearman correlation in frame-to-frame spike velocity and movement angle over the full AFM observation for each pair of spikes. The number of spike pairs with *p* < 0.01 is highlighted in green; the *r_s_* and *p* values for these are listed in the table below. (**j**) Relative frequencies of step angles measured between successive frames (unresolved frames excluded).

### Maturation-dependent nanomechanical properties of virions

To investigate the nanomechanical parameters of DENV, we next performed 3DFM on individual IM and WT DENV particles^44,45^. HS-AFM acquired immediately before 3DFM analysis corresponded well with the topographical map reconstructed using 3DFM data (Fig. 5a–c, Extended Data Fig. 5–6). For each force–distance curve at each pixel within the scan area, we calculated and mapped the YM, stiffness, and maximum indentation depth. All three nanomechanical parameters were heterogeneously distributed throughout virions. Notably, while YM distribution and indentation depth did seem to correspond, regions of high or low YM did not necessarily coincide with regions of high or low stiffness (Fig. 5a–c, Extended Data Fig. 5–6). This suggests that YM and stiffness are measuring different nanomechanical aspects of the virion. With YM fitting the initial contact region of the FD curve, here we consider it a measure of the resistance to perturbation of the viral protein coat, while stiffness, measuring the innermost linear region of the FD curve, measures the penetrability of the nucleocapsid core. Consistent with this interpretation, we found that the region of reduced YM (relative to the hard mica substrate) extended to the topographical edge of all particles (Extended Data Fig. 5c, 6c), while the region of reduced stiffness was truncated, particularly in IM (spiky) particles (Extended Data Fig. 5d, 6d). This suggests that stiffness is measuring the nucleocapsid core; the membrane and embedded proteins that extend laterally beyond the core do not prevent the cantilever tip from reaching the mica substrate. Based on the stiffness distribution, we categorized particles as either showing a central minimum (Fig. 5a), a central maximum (Fig. 5c), or a mix of the two (Fig. 5b). Overall, we found that IM particles were predominantly of the central maximum type, while WT particles were most often of the mixed maximum/minimum type (Fig. 6b). Notably, many IM particles showed peripheral arcs of low YM/stiffness and high indentation depth (Fig. 5c, Extended Data Fig. 5c–d), while similar regions in WT particles were more circular and nearer to the center (Fig. 5a–b, Extended Data Fig. 6c–d). This suggests that the structure of the DENV nucleocapsid core differs according to maturation state. Our full-particle 3DFM approach also enabled separate nanomechanical characterization of the smooth and spiky regions within single mosaic WT virions. By binary-masking particles based on their reconstructed topography, we compared the YM and stiffness distributions within the spiky and smooth regions of three favorably oriented WT virions (Fig. 5d–f, Extended Data Fig. 7). Although the values of YM and stiffness varied between particles, we consistently found that within each particle, YM and stiffness was higher in smooth regions compared to spiky regions. The increased YM suggests that the smooth dimeric M–E conformation provides increased resistance to perturbation compared to the spiky trimeric prM–E conformation, while the increase in stiffness suggests that the properties of the core underneath also differ.

**Figure 5.**
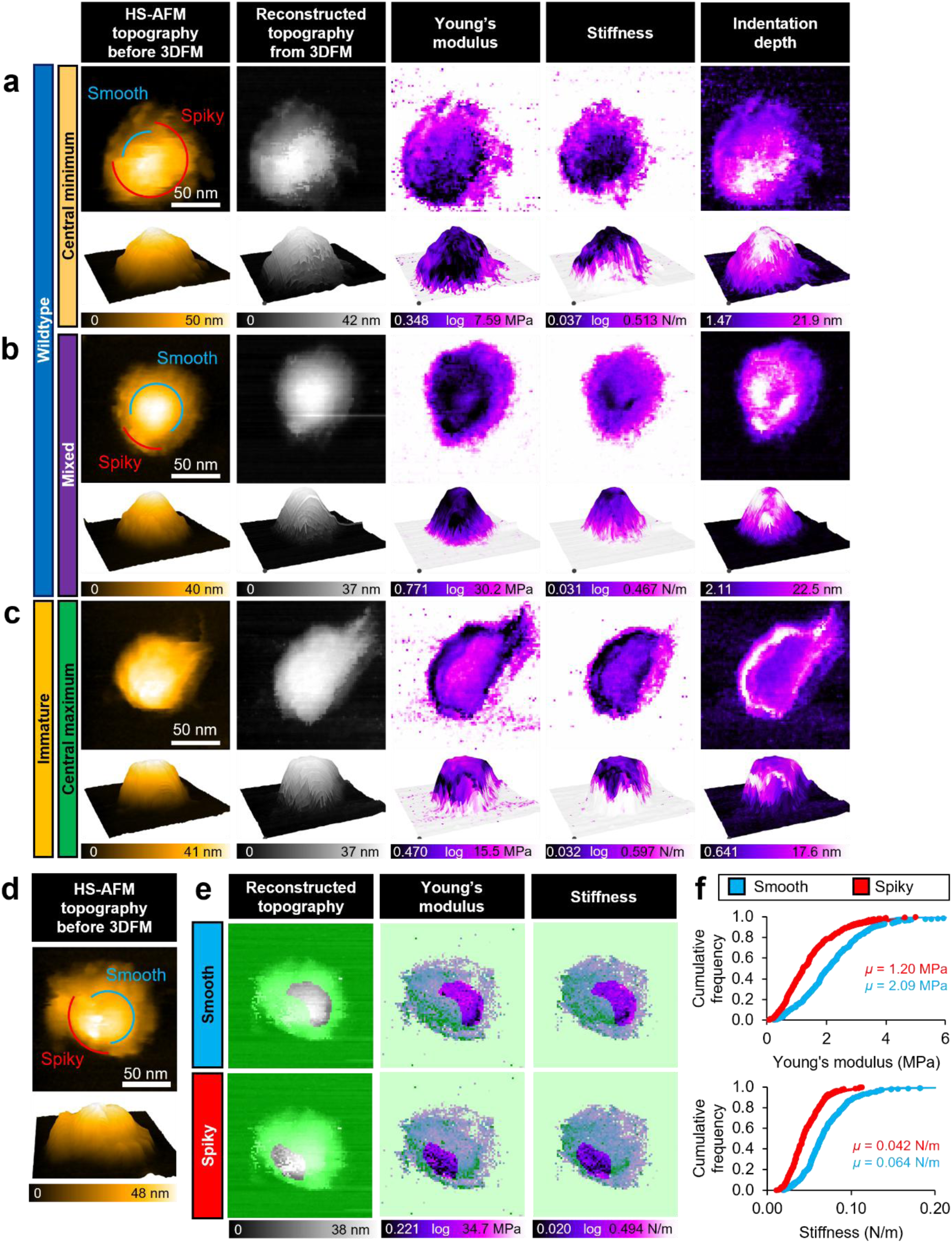
3DFM of DENV-2 virions reveals their nanomechanical properties at the sub-particle scale. Analysis of (**a**) WT particle showing central minimum stiffness; (**b**) WT particle showing a mix of central maximum and minimum stiffness; and (**c**) IM particle showing central maximum stiffness. (**a**–**c**) Left column: HS-AFM topography before 3DFM, acquired and processed as in Fig. 3. Right four columns: 80 × 80 × 180 pixels over 150 × 150 × 180 nm^3^ scan volume, scan speed 15.2 µm/s, frame time 1.6 s, scan time 121 s. Panel A maps are cropped to 80 × 78 × 180 pixels to remove an artefact at the bottom. Processing: median filter (*r* = 1 pixel). Scale bars: 50 nm. Lower panels show a 3D projection of the upper panel, with the Z axis exaggerated by 500% and smoothed by a Gaussian filter (*σ* = 2.0). (**d**) HS-AFM topography of a single WT virion. (**e**) Reconstructed topography, YM, and stiffness maps for the particle in (**d**). Each are masked (green shading) to show only the smooth region (upper panels) or spiky region (lower panels). (**f**) Cumulative distribution of YM (upper) and stiffness (lower) in the masked smooth and spiky images.

**Figure 6.**
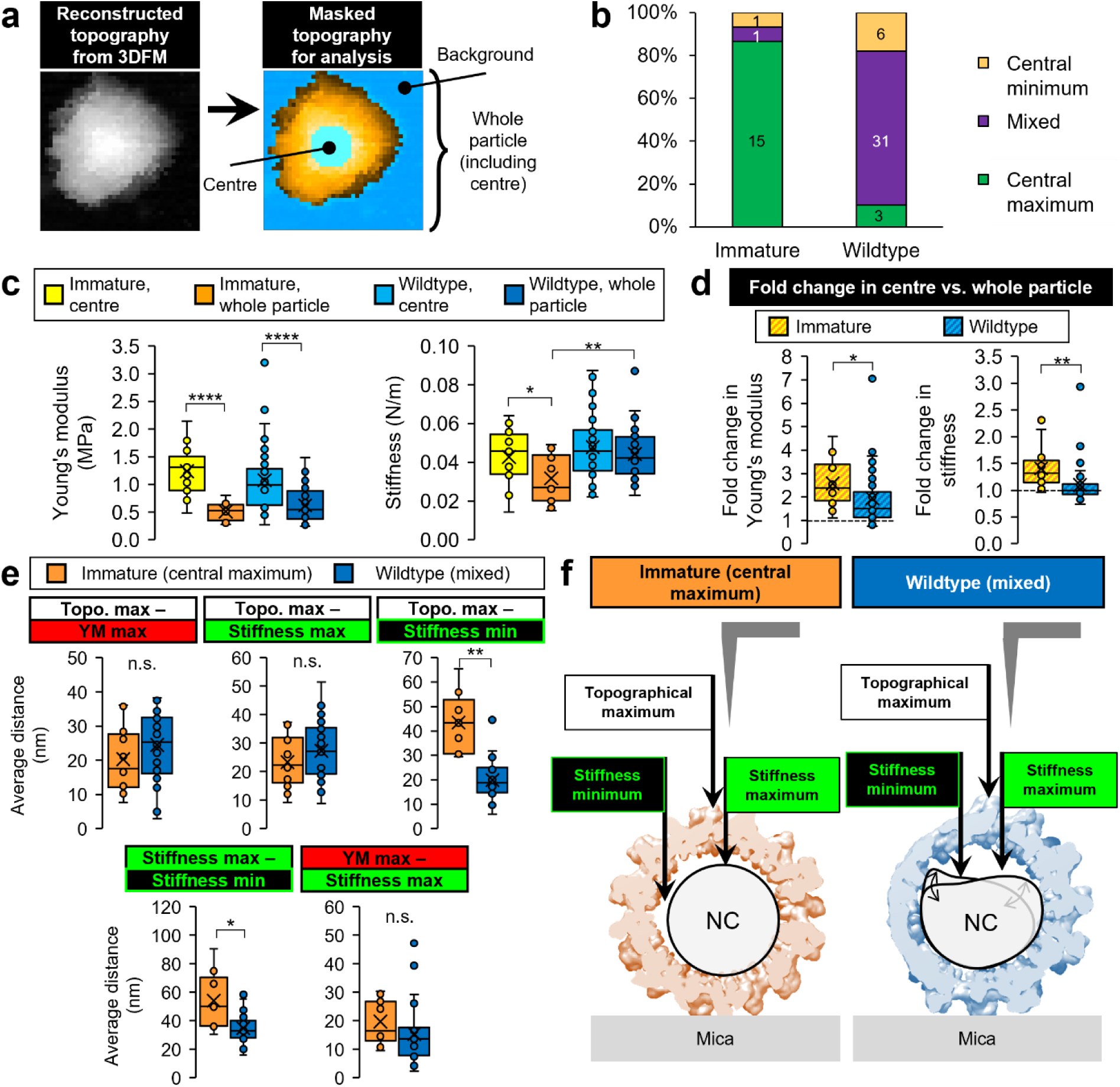
The nanomechanical properties of DENV-2 virions vary according to their maturation state. (**a**) Definition of regions used here. The ‘centre’ of the particle is a circular region comprising about 20% of the particle area, centred on the maximum height; the whole particle includes the centre but not background, thresholded by height using reconstructed topography. (**b**) Classification of each IM and WT particle characterized by 3DFM as showing a central nanomechanical minimum (green), maximum (yellow), or a mosaic of maxima and minima (purple). (**c**) Summary of YM (left) and stiffness (right), comparing the centres and whole particles for IM and WT virions. (**d**) For each individual particle, the fold change in Young’s modulus (left) and stiffness (right) in the centre compared to the whole particle. (**e**) The average 2D distances within individual particles among topographical, Young’s modulus, and stiffness maxima as well as stiffness minima. (**a**–**e**) Yellow: IM centre; orange: IM whole particle; light blue: WT centre; dark blue: WT whole particle. IM, *n* = 17; WT, *n* = 40. Boxes: interquartile range; line: median; cross, mean; error bars: SD.*, *p* < 0.05; **, *p* < 0.01; ****, *p* < 0.0001; n.s., not significant by Welch’s t-test. (**f**) Model of the differing internal properties between IM (central maximum) and WT (mosaic) virions. IM virions show a less stiff but less fluid nucleocapsid (NC) core, whereas WT virions show a stiffer but more fluid NC core, suggesting higher viscosity.

We next determined whether the nanomechanical differences observed at the sub-particle level were consistent across multiple virions. To this end, we performed 3DFM on IM (*n* = 17) and WT (*n* = 40) particles and calculated mean YM and stiffness values for each map, excluding the substrate and debris by thresholding the maps at 5–25% maximum height. To ensure like-to-like comparisons considering the geometry of the particles, we defined a ‘center’ region enclosing about 20% of the area of the particle, centered on the topographical maximum; the nanomechanical properties of this center were evaluated in comparison to the whole particle (including the center) (Fig. 6a).

Unsurprisingly, we found that YM was significantly higher in the center for both IM and WT particles (1.2 ± 0.4 and 1.1 ± 0.6 MPa respectively) compared to the entire particle (0.51 ± 0.16 and 0.62 ± 0.30 MPa respectively) , reflecting the increased ability of the protein coat to resist perpendicular (compressive) force over shearing force (Fig. 6c–d left). However, while IM stiffness was slightly increased in the center compared to the whole particle (0.043 ± 0.014 vs. 0.032 ± 0.012 N/m), WT stiffness was not significantly altered (0.048 ± 0.017 vs. 0.044 ± 0.013 N/m) (Fig. 6c–d right). This is consistent with the categorization that IM and WT particles show central nanomechanical maxima and mixed maxima/minima respectively. Compared to other human enveloped viruses, we found DENV-2 to be much less stiff compared to mature HIV-1 (0.22 N/m) or immature HIV-1 (3.15 N/m)^30^, but about threefold stiffer than SARS-CoV-2 (0.013 N/m), which lacks the organized protein capsid or coat seen in HIV or DENV respectively^46^. YM values were in line with those observed for Zika virus, another member of the *Flaviviridae* family, which were found to be around 0.2 to 0.9 MPa^47^.

Notably, whole-particle stiffness was significantly higher in WT than IM particles, agreeing with the trend seen within individual virions and further suggesting a maturation-dependent shift in core structure (Fig. 6c right). Interestingly, we found that particle maximum height was negatively correlated with stiffness for WT but not IM particles (Extended Data Fig. 8b). This contrasts with individual pixels within single virions, where height and stiffness were negatively correlated for WT as well as IM particles (Supplementary Fig. 6). We speculate that the more confined volume of lower-height particles results in greater internal pressure and thereby increased core resistance to perturbation; the effect of internal pressure on stiffness is well-characterized for lipid nanovesicles^48^, and similar effects have been reported for other viruses^26,43^. Interestingly, YM and stiffness were positively correlated for WT and IM particle centers, but not whole IM particles (Extended Data Fig. 8c), suggesting a maturation-dependent disconnect between peripheral and central nanomechanical resistance.

To better understand these maturation-dependent differences, we analyzed thresholded and smoothed YM and stiffness maps to determine the 2D distance among the topographical maximum, and maxima and minima of YM and stiffness, comparing IM (central maximum; *n* = 13) (Extended Data Fig. 5) and WT (mixed; *n* = 28) (Extended Data Fig. 6) particles. We found that significant YM minima were rarely observed for IM particles; we therefore excluded that parameter from our analysis. Among the remaining parameters, we found that the distance between topographical maximum (white dots) and minimum stiffness (green circles) was significantly increased in IM (central maximum) over WT (mixed) particles (43 ± 13 vs. 20 ± 8 nm respectively), and the distance between stiffness maxima (green dots) and minima (green circles) was similarly increased (54 ± 20 vs. 35 ± 11 nm respectively) (Fig. 6e, Extended Data Fig. 5e, 6e).

In light of these results and the peripheral arc-like pattern of stiffness minima seen in IM particles (Fig. 5c, Extended Data Fig. 5d), we propose a model whereby the core of IM particles is less fluid than WT particles, with a possibly gel-like nucleocapsid that does not occupy the entire membrane-enclosed volume of the virion (Fig. 6f left). Gaps between the off-center gel-like core and the viral membrane would thus give rise to the peripheral stiffness minima arcs we observed. In contrast, we propose that the core of WT particles is more fluid and under greater internal pressure, resulting in a higher overall stiffness value compared to IM as well as stiffness minima that are closer to the particle center as the core is pushed aside more easily by the invading cantilever tip (Fig. 6f right).

## Conclusions

The ensemble-averaging techniques that form the basis of our understanding of viral architecture preclude analysis of dynamic and disordered regions. Structural details and variations at the single-virion level are also lost, which leaves knowledge gaps in the context of conformationally diverse viruses such as DENV. Here, we applied HS-AFM and 3DFM to gain insight into the maturation-dependent changes in the dynamic structure and nanomechanical properties of DENV. While all virions show heterogeneity, PM (mosaic) DENV take on a particularly heterogeneous dynamic structure that is unlike the MA (smooth) or IM (spiky) conformations (Fig. 2–3). This emphasizes the usefulness of single-particle approaches to structural characterization, allowing observation of sub-virion phenomena that would be lost by ensemble averaging. Despite the overall stability of IM virions, we found that their prM–E spikes undergo confined independent movement, with variations in movement radius, velocity, and angle (Fig. 4). The mechanism underlying these dynamic differences among spikes in a single virion remains unclear. Incomplete or differential *N-*glycosylation across the virion surface may be a factor, as flaviviral E proteins are heavily glycosylated^49^ and *N-*glycosylation has been shown to reduce protein dynamics^50^. 3DFM of DENV further underlined the heterogeneity between and within DENV particles, with maturation-dependent changes in their nanomechanical properties and internal structure (Fig. 5–6). Our single-particle nanometrological approach could be readily applied to other enveloped viruses. Complementing existing viral structural techniques, this provides a foundation for investigating the real-time properties of individual virus particles in a biological-like context, adding another dimension to our understanding of viral biology.

## Acknowledgments

We thank Toshio Ando for his continuous support and encouragement. We also thank Kayo Nakatani and Risa Omura for their technical support.

## Funding sources

This research was funded, in part, by KAKENHI, Japan Society for the Promotion of Science [20H00327 and 24H00402 (to N.K.), 21K04849 (to K.U.), and PE20014 and 24K18450 (to S.J.M.)], AMED-CREST; the Japan Agency for Medical Research and Development [JP24gm1610009 (to N.K.)]; PRESTO, Japan Science and Technology Agency [JPMJPR20E3 and JPMJPR23J2 (to K.U.)]; and the World Premier International Research Center Initiative (WPI, MEXT, Japan, Kanazawa University).

## Author contributions

S.J.M. and N.K. designed the study. S.J.M. generated all samples, and designed and performed all experiments and analysis. K.U. designed and implemented the analysis software (UMEX Viewer and UMEX 3D Force Map Viewer). S.J.M., K.U., and N.K. wrote the paper, with contributions from all authors.

## Competing interests

The authors declare no competing interests.

## METHODS

### Cell lines and viruses

Vero cells (African green monkey kidney) were obtained from the JCRB Cell Bank (#9013) and maintained in minimum essential medium (MEM) supplemented with 10% fetal bovine serum (FBS), 1 mM sodium pyruvate, non-essential amino acids (NEAA), and 100 U/mL penicillin/streptomycin (Gibco/Thermo Fisher Scientific). DENV serotype 2, strain New Guinea C was obtained from ATCC (VR-1584) and propagated in Vero cells. Viral samples were created by inoculating 90% confluent Vero cell monolayers in 175 cm^2^ flasks at a multiplicity of infection (MOI) of 0.1 for 1 h, followed by incubation at 37°C with 5% CO_2_ in virus growth medium (VGM: MEM with 2% FBS and NEAA, yielding WT-DENV) or immature virus growth medium (IM-VGM: MEM with 2% FBS, NEAA, and 20 mM ammonium chloride, yielding IM-DENV^7,35^) with daily media changes until cytopathic effects became evident. Media (15 mL) were then harvested and clarified (3200 × *g*, 20 min, 4°C) to remove debris.

### Virus purification

Viruses were first concentrated by precipitation. PEG-6000 was added to the clarified infected cell culture supernatant (final concentration 8% w/v) for 18 h at 4°C, followed by pelleting (3200 × *g*, 8 h, 4°C). Virus-containing pellets were reconstituted in 0.9 mL PCSE buffer (phosphate-citrate-saline-EDTA: 20 mM sodium hydrogen phosphate, 10 mM sodium citrate, 100 mM sodium chloride, 1 mM EDTA, pH 7.4). Viruses were then purified by sucrose density gradient. 440 µL of each sample was loaded on a 4.4 mL 15–55% continuous sucrose gradient and ultracentrifuged (111,000 × *g*, 1.5 h, 4°C). Fractions of 400 µL were then collected, concentrated and exchanged into PCSE (final volume 40 µL) by ultrafiltration on 100 kDa MWCO centrifugal filter units (Millipore), and stored at –80°C.

### Plaque assay

DENV infectivity was quantified by semisolid overlay plaque assay in Vero cells. Vero cells at 90% confluence in 12-well plates were inoculated with 230 µL of sample and incubated at 37°C with 5% CO_2_ for 1 h, briefly rocking the plates at 20 min intervals to ensure wells did not dry out. The overlay medium (2% low melting point agarose (Takara), melted by autoclaving, in VGM) was kept liquid at 42°C. Following inoculation, 1.5 mL overlay was rapidly added to the infected cells, and the overlay was allowed to gel for 2 min at room temperature. Plates were then incubated for 5 days at 37°C with 5% CO_2_. Fixation was performed by adding 10% formalin in PBS directly to plates and incubating at RT for 1 h. Plates were then thoroughly rinsed with water, removing the agarose plugs, and stained for 5 min with crystal violet (0.1% w/v in 20% ethanol). Plaques were enumerated manually.

We confirmed the expected lack of IM-DENV infectivity by plaque assay in Vero cells, finding that infectivity was reduced below the limit of detection (6-log reduction) in IM samples compared to WT (Supplementary Fig. 1b–c). Virions in both samples were concentrated, purified, and exchanged into PCSE for HS-AFM observation. WT samples showed a peak infectivity of 1.8×10^8^ plaque-forming units (PFU) per mL. To render samples safe for HS-AFM observation was abolished by UV irradiation (*λ*_max_ = 253.7 nm) as previously described^51^, with 1 minute of exposure per log PFU/mL found to be sufficient to reduce infectivity below the limit of detection (Supplementary Fig. 2a–b).

### Simulated AFM

We used BioAFMViewer software^37^ to perform simulated AFM on the virion structure PDB files shown in Fig. 1 and Extended Data Fig. 1. Simulation conditions included a tip cone half angle of 8° and a tip apex radius of 5 nm that resembles the cantilevers we used for HS-AFM, with a scan step of 0.50 nm.

### HS-AFM sample preparation

Following inactivation, 2 µL of the sample was loaded onto a freshly cleaved mica disc (1.5 mm diameter, ∼0.05 mm thickness), which was mounted on a glass stage (2 mm height, 2 mm diameter) attached to the HS-AFM Z-scanner. After incubating for 10 minutes at room temperature to allow binding, the substrate was washed with 20 µL of PCSE. The scanner was then inverted and mounted on the HS-AFM, immersing the stage in the liquid cell containing 90 µL of PCSE for imaging.

### HS-AFM imaging

All imaging was performed on a laboratory-built HS-AFM operating in tapping mode^52^. Short cantilevers (9 × 2 × 0.13 µm; BL-AC10DS-A2, Olympus) were used, with a spring constant of ∼100 pN/nm, resonance frequency of ∼500 kHz, and quality factor of ∼1.5 in water. An amorphous carbon tip was fabricated by electron beam deposition on the apex of the cantilever’s bird’s beak, with an average length of ∼500 nm and a tip apex radius of ∼5 nm. The cantilevers were oscillated at a frequency slightly below their first resonance frequency to enhance the force sensitivity^53^. The free oscillation amplitude *A*_0_ was set to 3 nm, and the set-point amplitude was set to 0.9 × *A*_0_. While wide-area scans (1000 × 1000 nm^2^) were performed in trace-retrace (normal scanning) mode, most single-particle HS-AFM imaging was performed in only-trace imaging mode, whereby the tip tracks the surface topography during trace, but is lifted during retrace to enable faster movement to the next scan line^54^; this reduced the frame acquisition time for 150 × 150 nm^2^ scans from 2.3 s to 1.5 s. To minimize the error saturation at the edges of virus particles, dynamic PID feedback was used to control the tip-sample distance as described previously^55^.

### HS-AFM analysis

All image processing and analysis were performed in a laboratory-built software, UMEX Viewer. Tilt correction in the XY plane was applied manually. Images were smoothed (Gaussian filter *σ* = 1) and denoised (median filter *r* = 1). Particle heights were calculated as the difference between the maximum particle height and the average height of a bare substrate region near the particle. In some images, frame averaging was performed over three consecutive frames using a triangular kernel to further reduce noise. For supplementary videos, translational drift correction was performed using the StackReg plugin^56^ in ImageJ. For roughness calculations, each frame (Gaussian smoothed but otherwise unfiltered) was processed by second-order flattening filters along both the X and Y directions, and a first-order flattening filter in the XY plane. The image was then cropped to a region corresponding to the upper surface of the virion (45 × 45 pixels = 45 × 45 nm^2^ for WT (smooth and mosaic) particles, 60 × 60 pixels = 60 × 60 nm^2^ for IM (spiky) particles), and the roughness value was calculated as the standard deviation of a Gaussian curve fit to the histogram of height deviations of each pixel relative to the flattened plane.

### Analysis of spike dynamics

HS-AFM image stacks (∼100 frames) were preprocessed by filtering (Gaussian *σ* = 1, median *r* = 1) and flattening (second order in X and Y, first order in XY plane) in UMEX Viewer, then exported at 512 × 512 pixels and imported into ImageJ, where translational drift correction was performed using StackReg. Individual spikes were tracked using the TrackMate plugin^57^ for ImageJ. Briefly, an elliptical ROI was defined for each spike. Spike peaks were identified by a Laplacian of Gaussian (LoG) detector (radius 20 pixels), and tracked with a Simple Linear Assignment Problem (LAP) tracker. Tracks were then manually curated to obtain the final time series, with gaps manually confirmed where the spike could not be localized.

The 2D distance between neighboring spikes in the cryoEM structure was calculated by performing simulated AFM as described above, preprocessing the simulated AFM image as described above and localizing the maximum using a lab-built software, UMEX Viewer for Height Analysis. Displacements were calculated relative to the average location of each spike over the time course. Frame-to-frame movement angle and velocity were calculated by comparing contiguous pairs of frames; frames in which the spike could not be tracked were excluded. Correlations between frame-to-frame velocity and angle among pairs of spikes were evaluated by calculating the Spearman correlation coefficient *r_s_*; this value was then used to calculate a *T* score and two-tailed *p* value.

Radius of confinement was calculated using the mean-square displacement (MSD) method^58^. Briefly, the MSD is a function of the frame lag *m* between different spike positions, defined as:

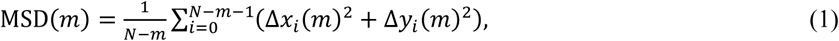

where *N* is the total number of frames and Δ*x_i_* and Δ*y_i_* are the X and Y displacements in the spike position between frames *i* and (*i + m*). In the case of constrained diffusion inside a circular region with radius of confinement *R_c_*, after correcting for localization error, MSD eventually plateaus at *R_c_^2^* (Supplementary Fig. 4). This gives an approximation:

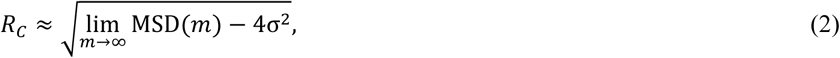

where *σ* is the localization error, estimated by the intercept *b* = 4*σ*^2^ of a linear model of the initial linear portion of the MSD curve:

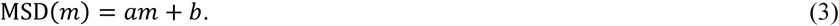

*R_c_* was calculated using *m* = [14, 25], with the linear fit over *m* = [1, 10] used to calculate *b* (Supplementary Fig. 4).

### 3D force mapping

All 3DFM was performed during HS-AFM experiments, with HS-AFM topography acquisition before and after force mapping. For each experiment, after retracting the cantilever from the substrate, Brownian noise fitting was performed to approximate the spring constant of the cantilever and calculate the inverted optical laser deflection detection sensitivity. Then, after centering on the particle of interest in HS-AFM tapping mode, the feedback signal was held constant and cantilever oscillation stopped to allow 3DFM. The probe was indented into the sample at a constant tip speed of 15.2 µm/s; retraction was triggered by a deflection signal of 0.1 ∼ 0.2 V, which was set during mapping to correspond to an apparent maximum loading force of ∼1.0 nN (∼0.6 nN after data processing).

### 3D force map analysis

All data processing and analysis was performed in a laboratory-built software, UMEX 3D Force Map Viewer. Tilt correction in the XY plane was applied manually, with the flattening threshold set to ∼50% of maximum deflection. Baseline correction was performed for each force curve by fitting and subtracting the background in the far region of the force curve: a 0th-order polynomial over the first 10% of Z-piezo extension, and a first- or second-order polynomial over the remaining background region until the tip begins to interact with the sample. Exact values for the inverted optical lever sensitivity (invOLS) were determined by calibrating force curves on the bare mica substrate and were typically in the range of 15−25 nm/V. For subsequent mechanical analysis, the Z-piezo extension for each force curve was converted to the tip–sample separation distance (Z-distance) using invOLS. To perform quantitative force curve measurements, it is important to consider the dynamic-to-static correction factors for both the spring constant and invOLS. However, in our setup, the cantilever length and the laser spot size are both approximately 9 μm, with the laser positioned near the center of the cantilever^53^. Therefore, as each of these correction factors affects the results by only about 3%—which is sufficiently smaller than the measurement error—no correction was applied.

Young’s modulus (YM) of the virus, *E*_virus_, was determined for each force curve by fitting with the Hertzian model as follows:

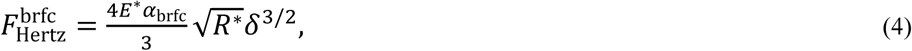

where *δ* denotes the indentation depth. *R*\* denotes the reduced radius, which is given by

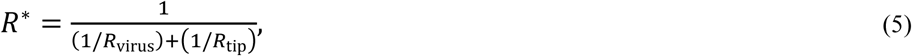

where *R*_virus_ is the radius of virus particle, determined from the particle height measured by HS-AFM immediately before 3DFM, and *R*_tip_ is the radius of the tip, assumed to be 10 nm. *E*\* denotes the reduced YM, which is given by

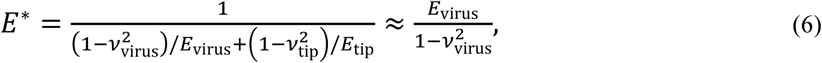

where the YM of the tip, *E*_tip_, was assumed to be sufficiently larger than *E*_virus_. *ν*_tip_ and *ν*_virus_ denote the Poisson’s ratio of the tip and virus, respectively, and a value of 0.5 was assumed for *ν*_virus_.

The Garcia bottom effect correction was also applied to accounts for the apparent overestimation of YM due to the reactive force from a rigid substrate^59^, whose factor, *α*_brfc_ is given by

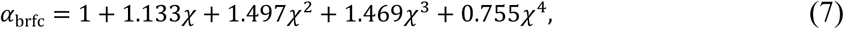

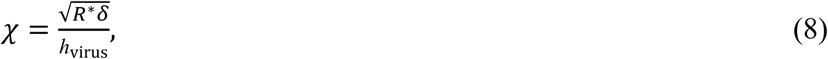

where *h*_virus_ is the virus height, given by 2×*R*_virus_. Although originally developed for flat membranes, it provides a reasonable approximation for spherical particles larger than the tip radius, such as viruses. Another bottom effect in spherical particles involves deformation of the particle’s bottom surface, which can lead to an underestimation of YM^60^. For ∼50 nm particles, this effect may reduce the modulus by up to half under the assumption of a perfect sphere. However, due to surface adhesion, the bottom surface is reasonably assumed to be flattened even in the absence of tip contact^61^, likely limiting the extent of this effect. Taken together, these two bottom effects are thought to largely cancel each other out, resulting in a net deviation from the Hertz model of less than ∼20% (Supplementary Fig. 5).

Stiffness was determined for each force curve by calculating the linear slope of the force curve after the tip passed through the Hertz fitting region. Whole-particle statistics were determined by setting a height threshold at 5–20% of the maximum height to delimit the particle and exclude debris, followed by lognormal fitting on the frequency distribution of YM and stiffness. Although no smoothing was applied before calculating nanomechanical parameters for each map, a Gaussian filter (*σ* = 1.0) was applied to the maps shown in figures to reduce noise.

Localization of maxima and minima for reconstructed topography, YM, and stiffness (Extended Data Fig. 5–6) was performed in ImageJ. Briefly, 8-bit force maps were preprocessed by a median filter to reduce high-frequency noise (*r* = 1 for 40 × 40 pixel maps, *r* = 2 for 80 × 80 or 90 × 90 pixel maps), upscaled to 512 × 512 pixels with bicubic interpolation, and smoothed by a Gaussian filter (*σ* = 7.0). Maps were globally thresholded on the reconstructed topography map using the triangle method^62^ and a binary mask was created to exclude everything but the virion. Maxima and minima were determined by calculating the centroid of all maximum or minimum pixels in the masked image, either globally or within a manually chosen ROI.

## EXTENDED DATA FIGURES

**Extended Data Figure 1.**
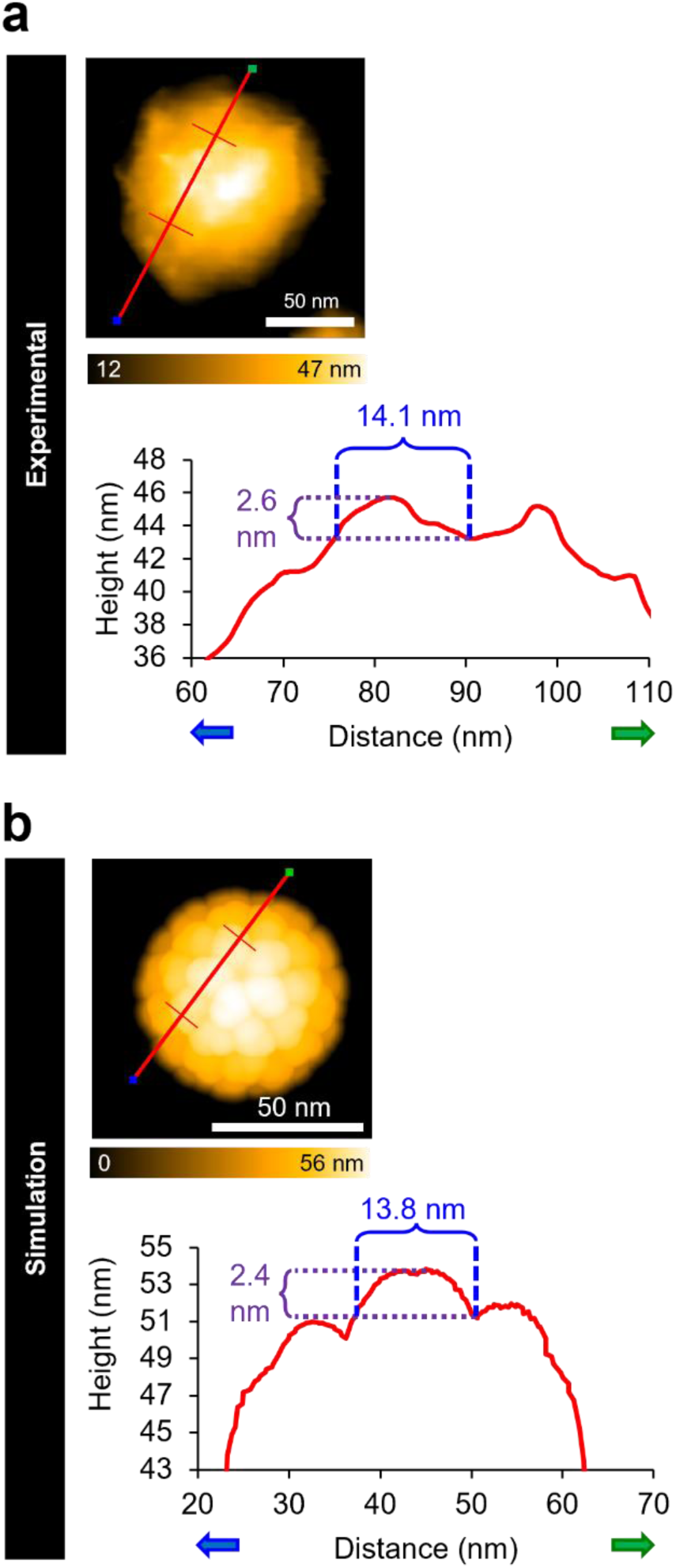
IM-DENV virion structures observed by HS-AFM resemble predictions based on existing cryoEM structures. (**a**) Representative HS-AFM image of an IM-DENV particle (scan area 150 × 150 nm^2^ at 150 × 150 pixels, scan speed 20 µm/s, frame time 1.5 s in only-trace mode). (**b**) Simulated AFM image of the IM-DENV cryoEM structure (PDB 3C6E), using a scan step of 1.0 nm, tip radius of 5.0 nm, and cone half-angle of 8°. (**a**–**b**) The spike-to-spike height profile along the red line from blue end to green end (cropped at the transverse red lines) is shown, measuring the approximate base diameter (blue dashed lines) and peak-to-trough prominence (purple dashed lines) of a single spike.

**Extended Data Figure 2.**
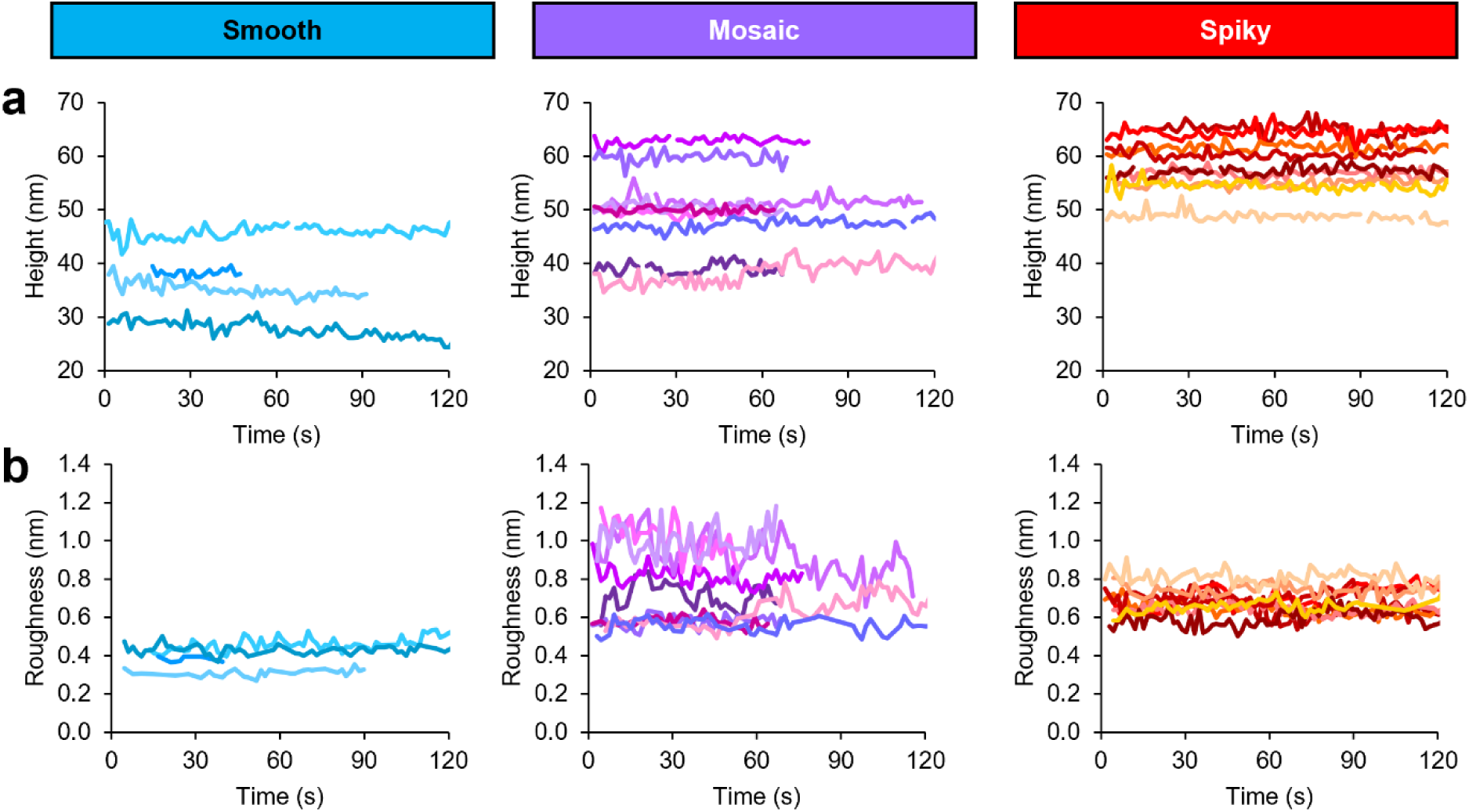
Single-particle observations show dynamic variations in height and roughness over time. (**a**–**b**) Maximum height (**a**) and surface roughness (**b**) measured by HS-AFM frame-by-frame for representative particles of each morphotype. Traces are distinguished by arbitrary colors. Smooth, *n* = 4; mosaic, *n* = 9; spiky, *n* = 9. Gaps are shown where HS-AFM imaging was unstable or abrogated by artefacts.

**Extended Data Figure 3.**
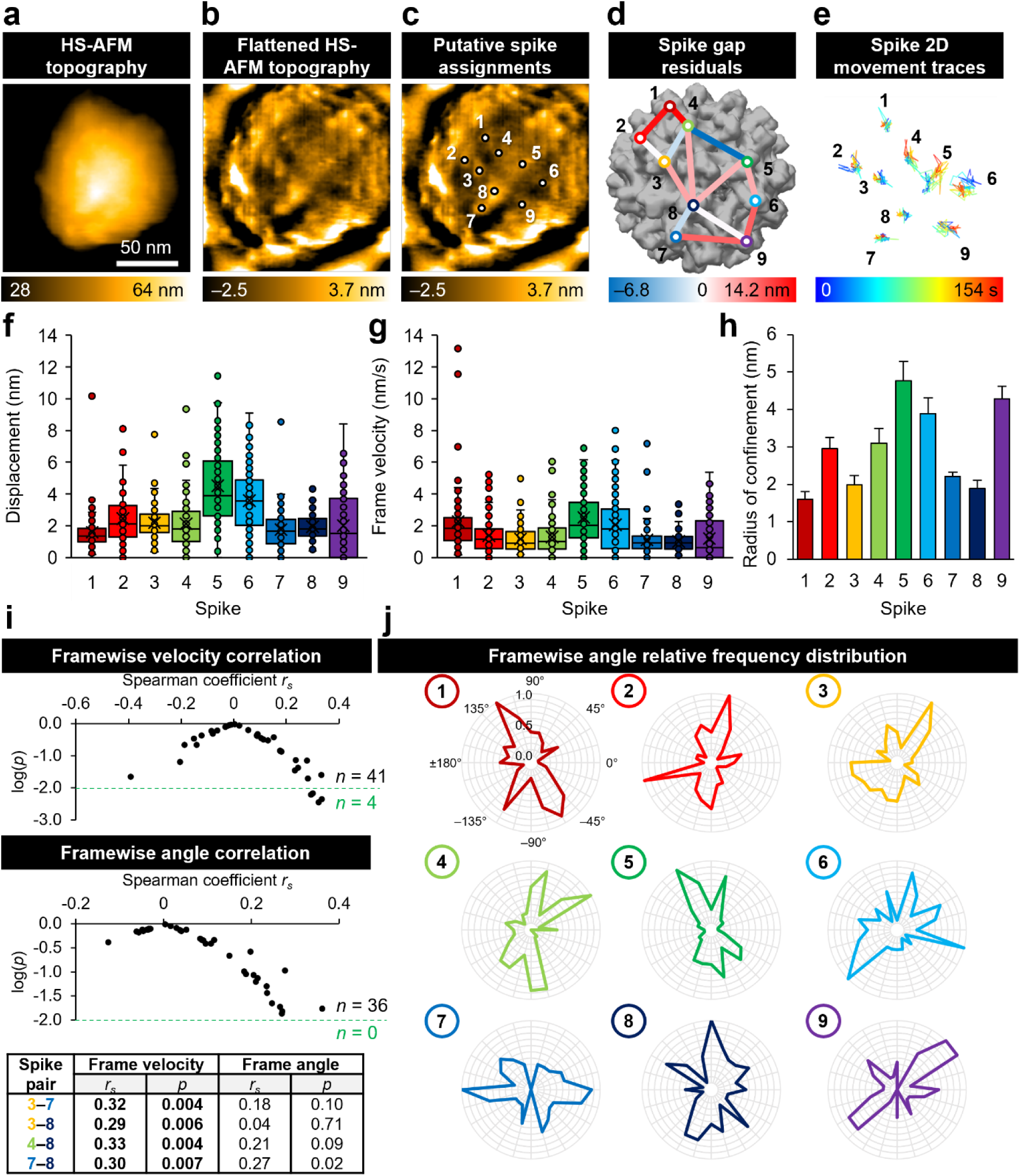
An additional example showing heterogeneous spike dynamics in IM-DENV virions. (**a**) Representative HS-AFM image (acquired and processed as described in Fig. 3; scale bar 50 nm). (**b**) The same image, smoothed and flattened. (**c**) The same image, with trackable spikes labeled and arbitrarily numbered. (**d**) Trackable spikes mapped onto the cryoEM structure (PDB 4B03). Residuals in the 3D distance between adjacent spikes in the cryoEM structure and HS-AFM structure are indicated. (**e**) Spike localization traces, coloured blue to red over time. (**f**) Displacement of each spike at each frame relative to the average position of the spike (unresolved frames excluded). (**g**) Spike velocity measured between successive frames (unresolved frames excluded). (**f**–**g**) Boxes: interquartile range; lines: median; crosses, mean; error bars: SD. (**h**) 2D apparent radius of confinement for each spike, calculated using the mean square displacement method. Error bars: SD. (**i**) Pairwise Spearman correlation in frame-to-frame spike velocity and movement angle over the full AFM observation for each pair of spikes. The number of spike pairs with *p* < 0.01 is highlighted in green; the *ρ* and *p* values for these are listed in the table below. (**j**) Relative frequencies of step angles measured between successive frames (unresolved frames excluded).

**Extended Data Figure 4.**
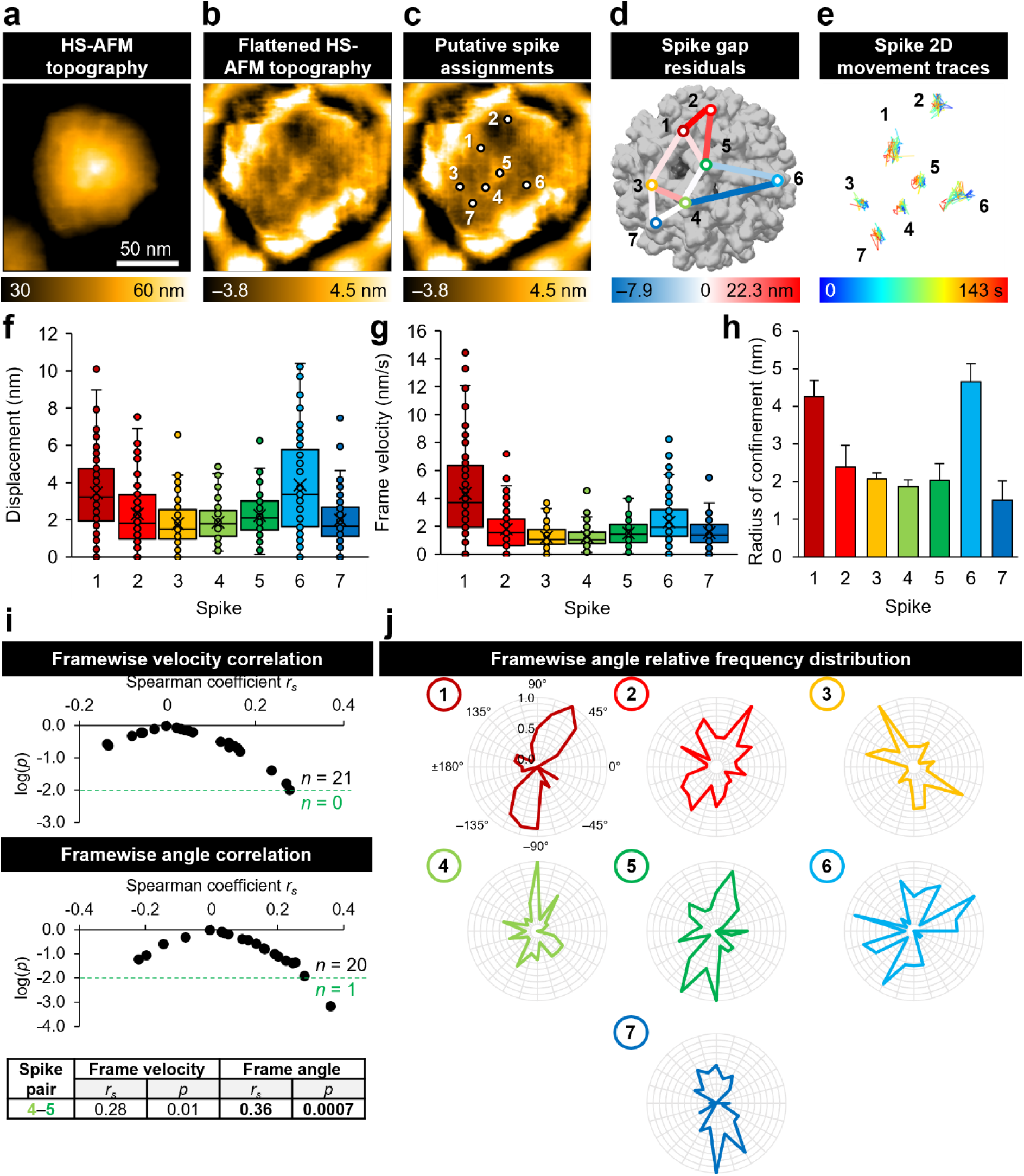
Another example showing heterogeneous spike dynamics in IM-DENV virions. (**a**) Representative HS-AFM image (acquired and processed as described in Fig. 3; scale bar 50 nm). (**b**) The same image, smoothed and flattened. (**c**) The same image, with trackable spikes labeled and arbitrarily numbered. (**d**) Trackable spikes mapped onto the cryoEM structure (PDB 4B03). Residuals in the 3D distance between adjacent spikes in the cryoEM structure and HS-AFM structure are indicated. (**e**) Spike localization traces, coloured blue to red over time. (**f**) Displacement of each spike at each frame relative to the average position of the spike (unresolved frames excluded). (**g**) Spike velocity measured between successive frames (unresolved frames excluded). (F–G) Boxes: interquartile range; lines: median; crosses, mean; error bars: SD. (**h**) 2D apparent radius of confinement for each spike, calculated using the mean square displacement method. Error bars: SD. (**i**) Pairwise Spearman correlation in frame-to-frame spike velocity and movement angle over the full AFM observation for each pair of spikes. The number of spike pairs with *p* < 0.01 is highlighted in green; the *ρ* and *p* values for these are listed in the table below. (**j**) Relative frequencies of step angles measured between successive frames (unresolved frames excluded).

**Extended Data Figure 5.**
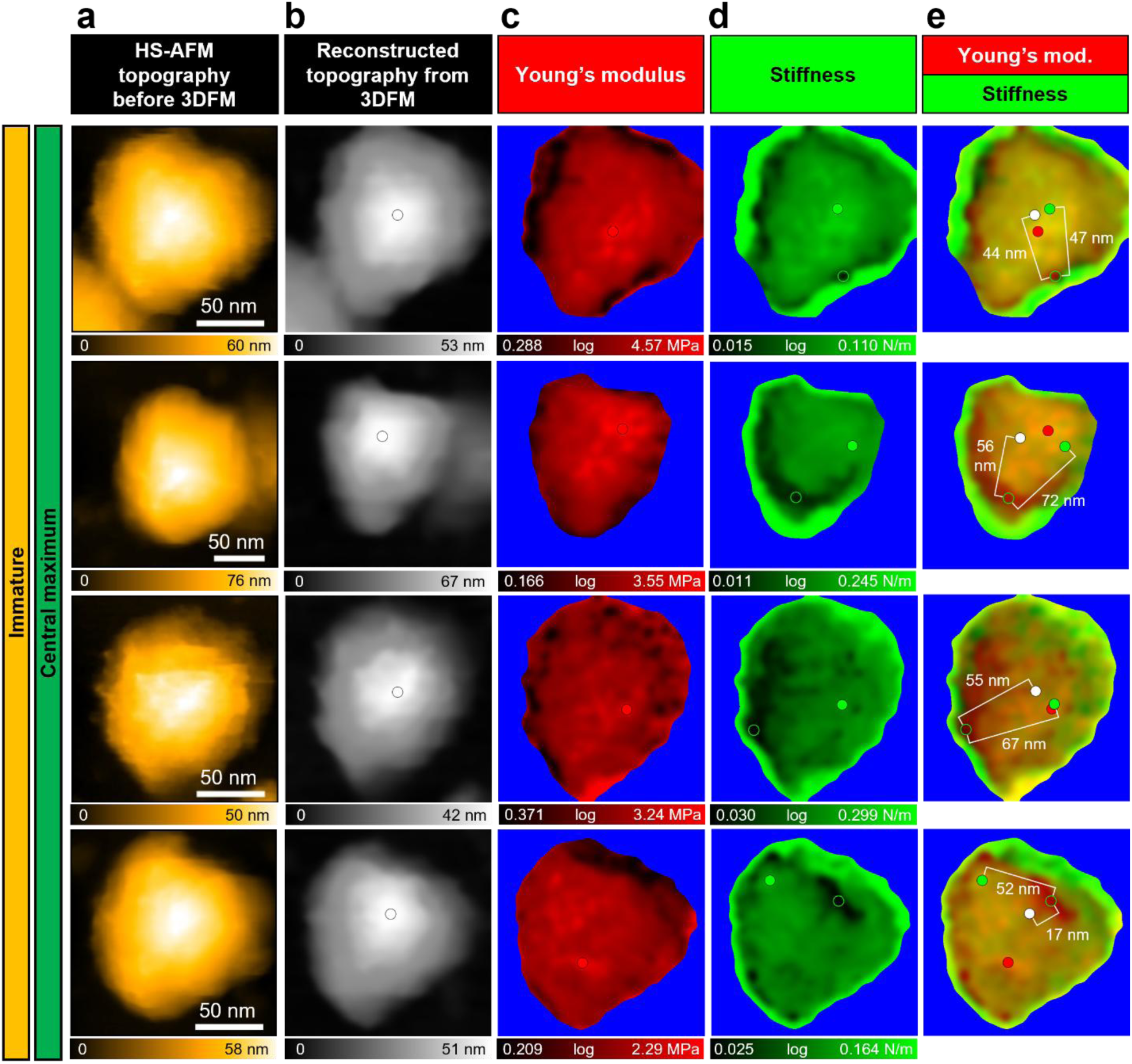
Additional wildtype virions showing sub-particle nanomechanical variations by 3DFM. (**a**) HS-AFM topography before 3DFM, acquired and processed as in Fig. 3. (**b**) Topography reconstructed from 3DFM data; white dot, maximum height. (**c**) YM map; red dot and circle, maximum and minimum YM respectively. (**d**) Stiffness map; green dot and circle, maximum and minimum stiffness respectively. (**e**) Merged YM and stiffness maps with corresponding maxima; color scales are the same as in panels C and D. Distances between stiffness minima and topographical/stiffness maxima are noted. (**B**–**E**) Maps acquired as 80 × 80 × 180 pixels over 150 × 150 × 180 nm^3^ scan volume, scan speed 15.6 µm/s, frame time 1.4 s, scan time 113 s (top and bottom row); 90× 88 × 180 pixels over 150 × 147 × 180 nm^3^ scan volume, scan speed 15.1 µm/s, frame time 1.6 s, scan time 140 s (second row); 90 × 90 × 180 pixels over 200 × 200 × 180 nm^3^ scan volume, scan speed 15.1 µm/s, frame time 1.6 s, scan time 148 s (third row). Processing: median filter (*r* = 2 pixels), Gaussian filter (*σ* = 0.5 pixels), thresholded by height using reconstructed topography. Scale bars: 50 nm.

**Extended Data Figure 6.**
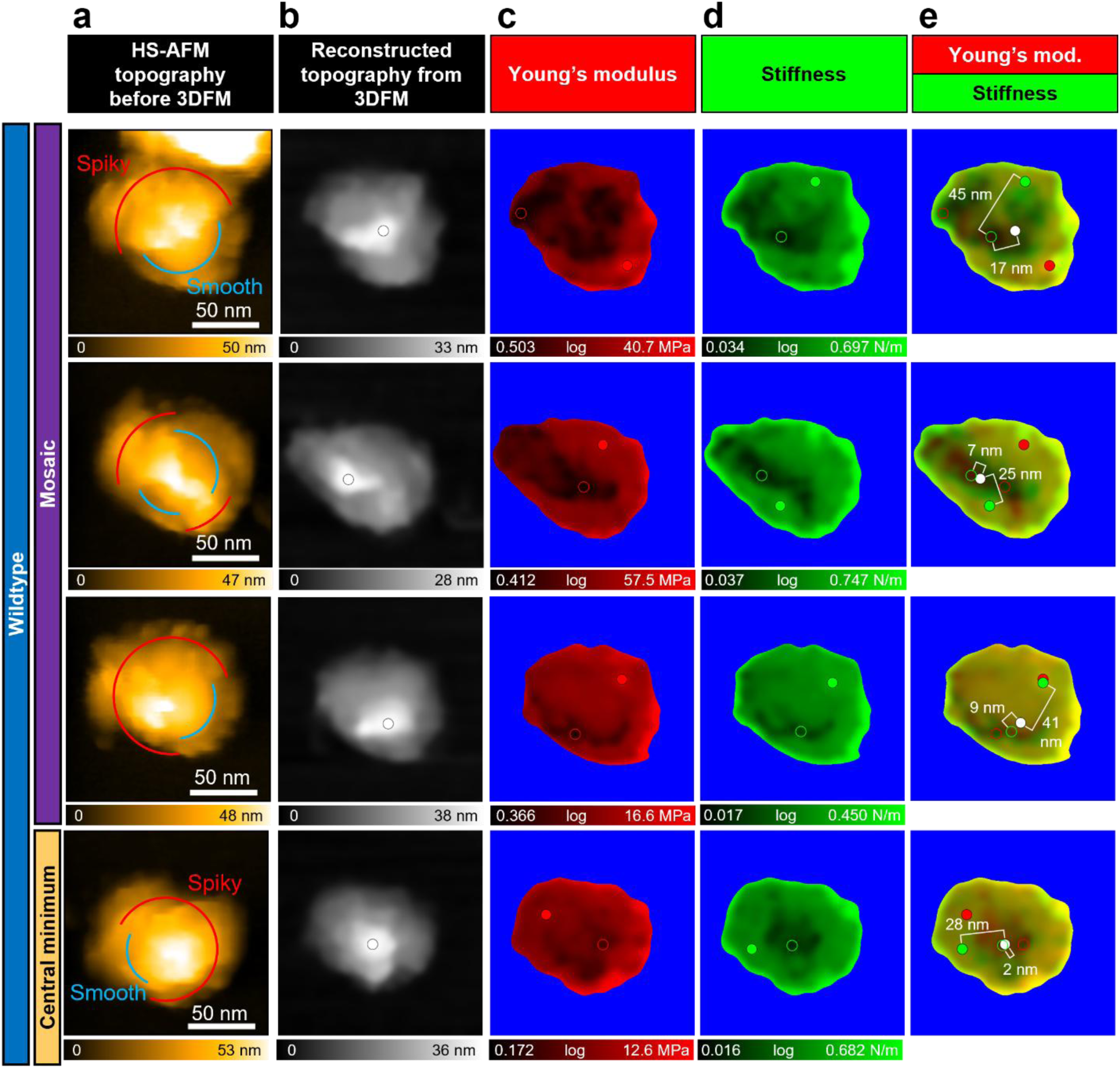
Additional immature virions showing sub-particle nanomechanical variations by 3DFM. (**a**) HS-AFM topography before 3DFM, acquired and processed as in Fig. 3. (**b**) Topography reconstructed from 3DFM data; white dot, maximum height. (**c**) YM map; red dot and circle, maximum and minimum YM respectively. (**d**) Stiffness map; green dot and circle, maximum and minimum stiffness respectively. (**e**) Merged YM and stiffness maps with corresponding maxima; color scales are the same as in panels C and D. Distances between stiffness minima and topographical/stiffness maxima are noted. (**b**–**e**) Maps acquired as 80 × 80 × 180 pixels over 150 × 150 × 180 nm^3^ scan volume, scan speed 15.3 µm/s, frame time 1.5 s, scan time 123 s (top row), 80 × 80 × 180 pixels over 150 × 150 × 180 nm^3^ scan volume, scan speed 15.2 µm/s, frame time 1.6 s, scan time 127 s (middle row); 80 × 80 × 180 pixels over 150 × 150 × 180 nm^3^ scan volume, scan speed 15.2 µm/s, frame time 1.6 s, scan time 125 s (bottom row). Processing: median filter (*r* = 2 pixels), Gaussian filter (*σ* = 0.5 pixels), thresholded by height using reconstructed topography. Scale bars: 50 nm.

**Extended Data Figure 7.**
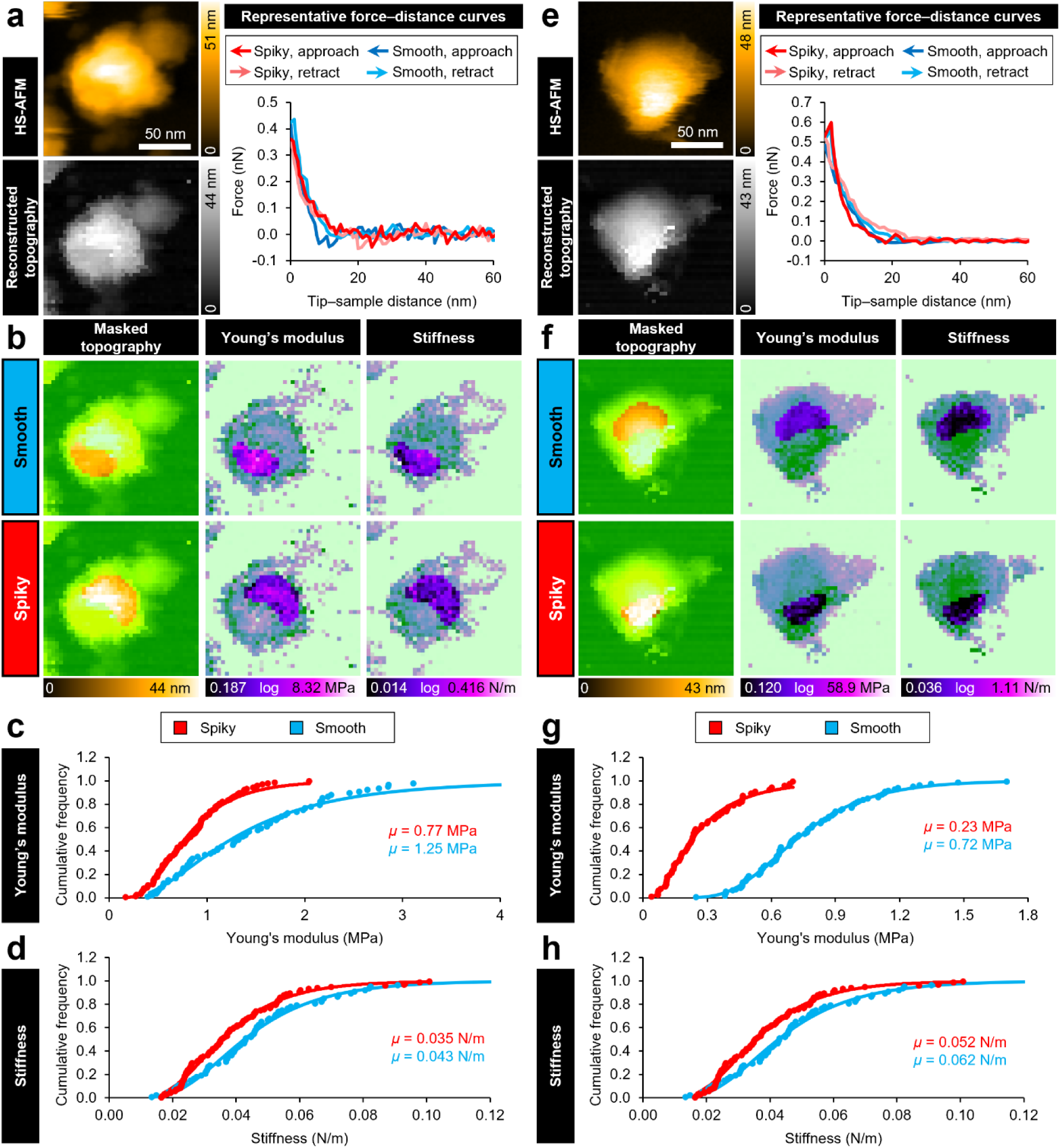
The spiky and smooth regions within mosaic virions show distinct nanomechanical properties. Two wildtype virions with mosaic morphology (**a**–**d** and **e**–**h**). (**a**, **d**) HS-AFM and reconstructed topography (left). Force-distance curves (right) for a representative pixel within the spiky/smooth regions shown in (**b**). The lowest 60 nm of the approach (dark red/blue, spiky/smooth respectively) and retract (light red/blue, spiky/smooth respectively) curves are shown. Scale bars: 50 nm. (**b**, **e**) Masked reconstructed topography, YM, and stiffness maps for the particles in (**a**, **d**). (**c**–**d**, **f**–**g**) Cumulative distribution for (**c**, **f**)Young’s modulus and (**d**, **g**) stiffness for all pixels in the spiky (red) and smooth (blue) regions. (**a**–**g**) HS-AFM images were acquired and analyzed as described in Fig. 3. Maps were acquired as described in Fig. 5 but not filtered or processed aside from masking.

**Extended Data Figure 8.**
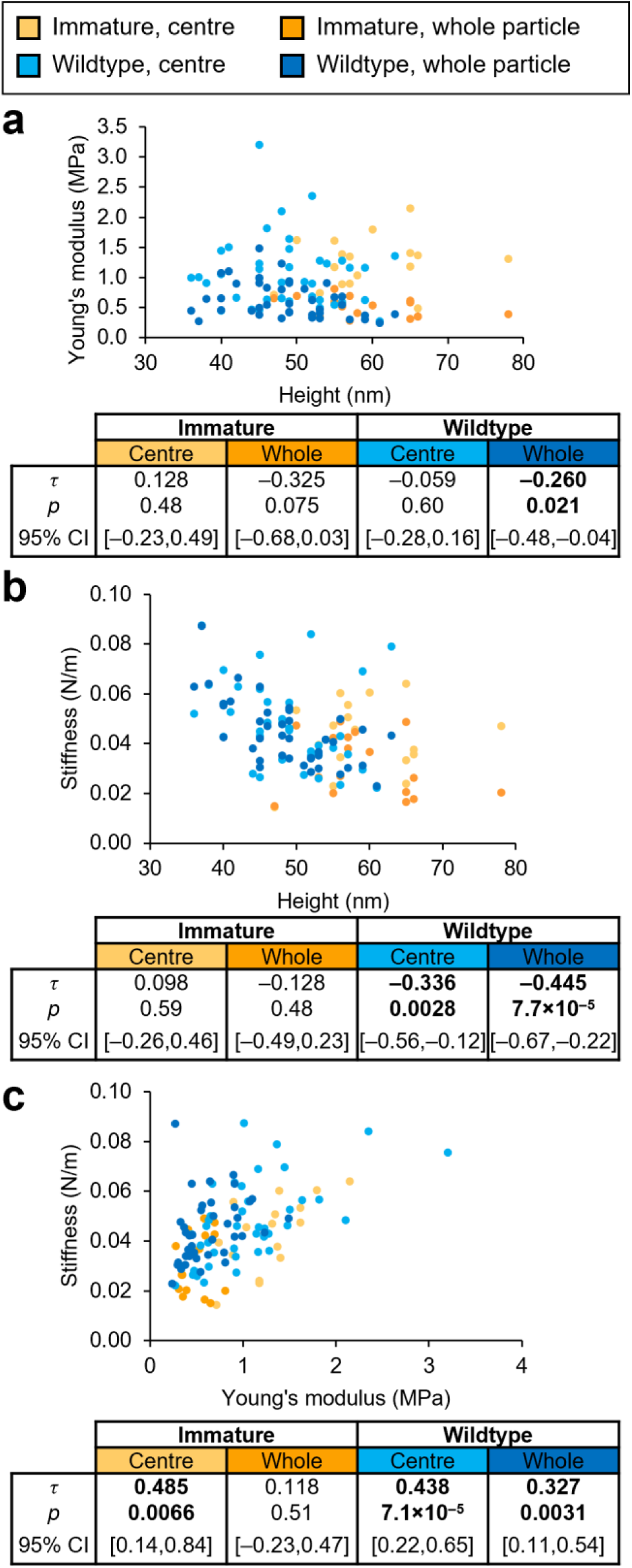
Height dependence and interdependence of nanomechanical parameters. Replotting of the (**a**) Young’s modulus and (**b**) stiffness data summarized in Fig. 6c as a function of virion height, as well as (**c**) stiffness as a function of Young’s modulus. Each point represents data for one virion, comprising the average values from 1–4 separate force maps. Yellow: IM centre; orange: IM whole particle; light blue: WT centre; dark blue: WT whole particle. IM, *n* = 17; WT, *n* = 40. *τ* is the Kendall rank correlation coefficient, used to calculate the two-tailed *p* value shown. Correlations (*p* < 0.05) are highlighted in bold.

## Notes

### Competing Interest Statement

The authors have declared no competing interest.

