## Supplementary Information for "Maturation-dependent structural dynamics and nanomechanical properties of dengue virus revealed by high-speed AFM and 3D force mapping"

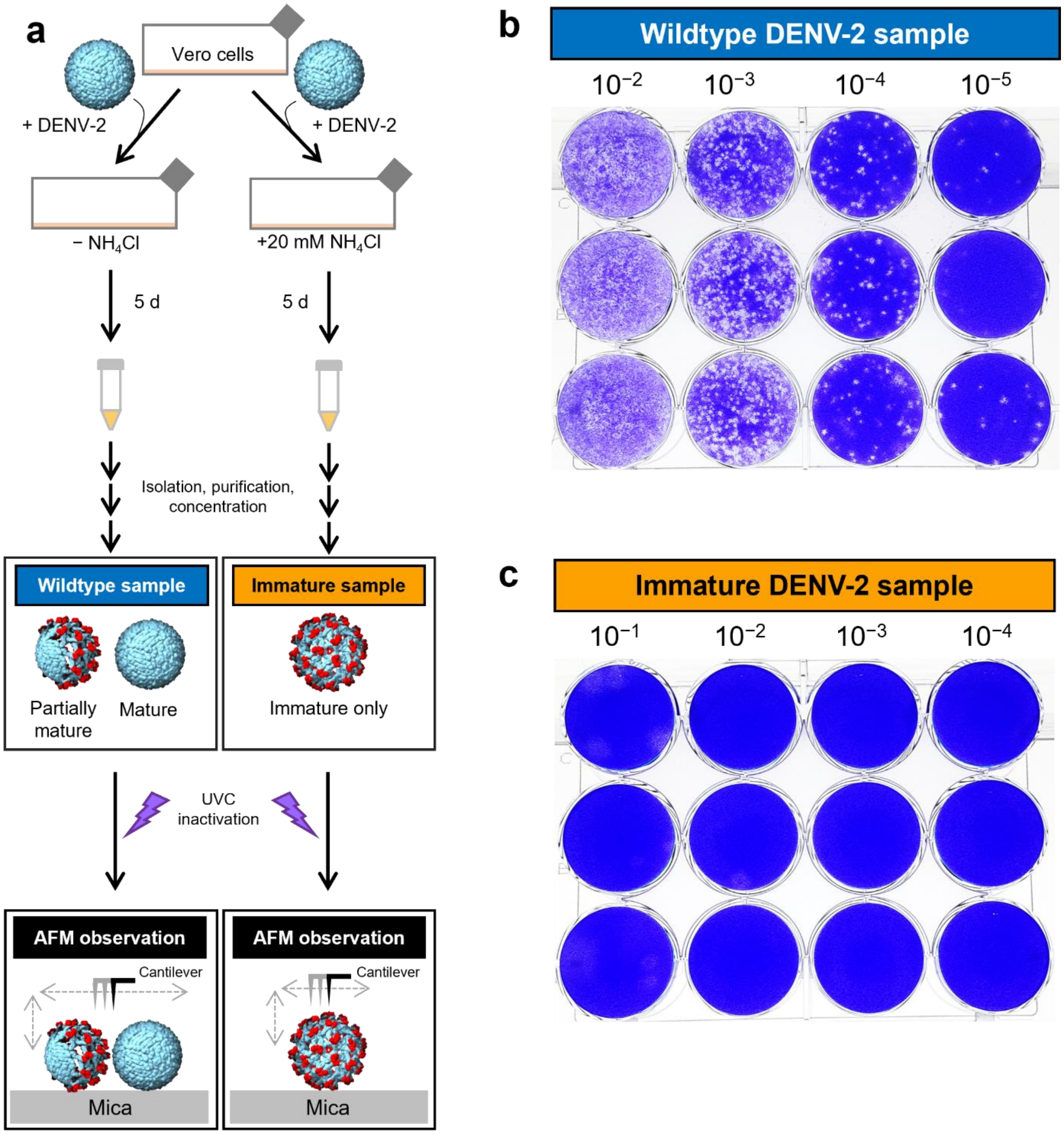


**Supplementary Figure 1. Immature DENV-2 preparations are non-infectious.** (**a**) Vero cells infected with DENV-2 produce a mixture of mature and partially mature virions ('wildtype'). Adding ammonium chloride to the culture medium yields fully immature virions[1]. Purified virions were inactivated by UV irradiation[2], bound to freshly cleaved mica, and imaged by HS-AFM in a liquid buffer (pH 7.4) at room temperature. (**b**–**c**) Representative plaque assays in Vero cells for DENV-2 grown in the absence (**b**) and presence (**c**) of 20 mM ammonium chloride, using non-purified cell culture supernatant as the inoculum in triplicate (columns) at the dilutions indicated (rows). Infectivity was reduced below the limit of detection in immature preparations.

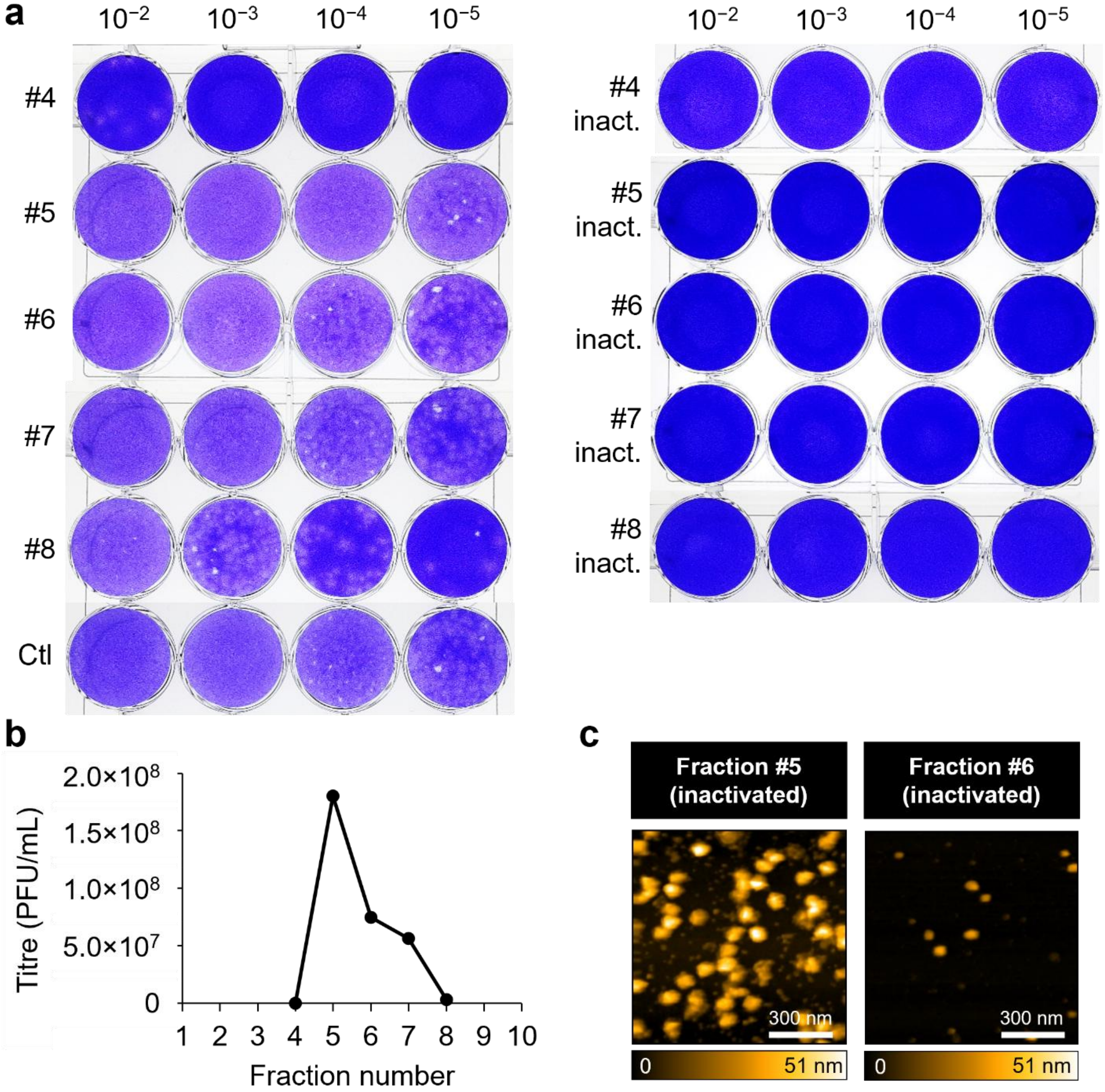


**Supplementary Figure 2. Characterization of purified WT DENV-2 infectivity and susceptibility to inactivation.** (**a**) Plaque assays measuring the infectivity of DENV-2 following purification, performed at the inoculum dilutions indicated. Numbers are sucrose gradient fractions. Ctl: original DENV-2 stock. Right panel: infectivity was abolished in samples inactivated by UV irradiation ('inact.'). (**b**) WT DENV-2 reached a maximum titre of $1.8 \times 10^8$ plaque-forming units (PFU) per mL in fraction #5. (**c**) Representative HS-AFM (scan area $1000 \times 1000$ nm$^2$, $200 \times 200$ pixels, scan speed 40 µm/s, frame time 10.3 s in trace-retrace mode) of fractions #5 and #6, showing virus-sized particles in #5 that are reduced in #6.

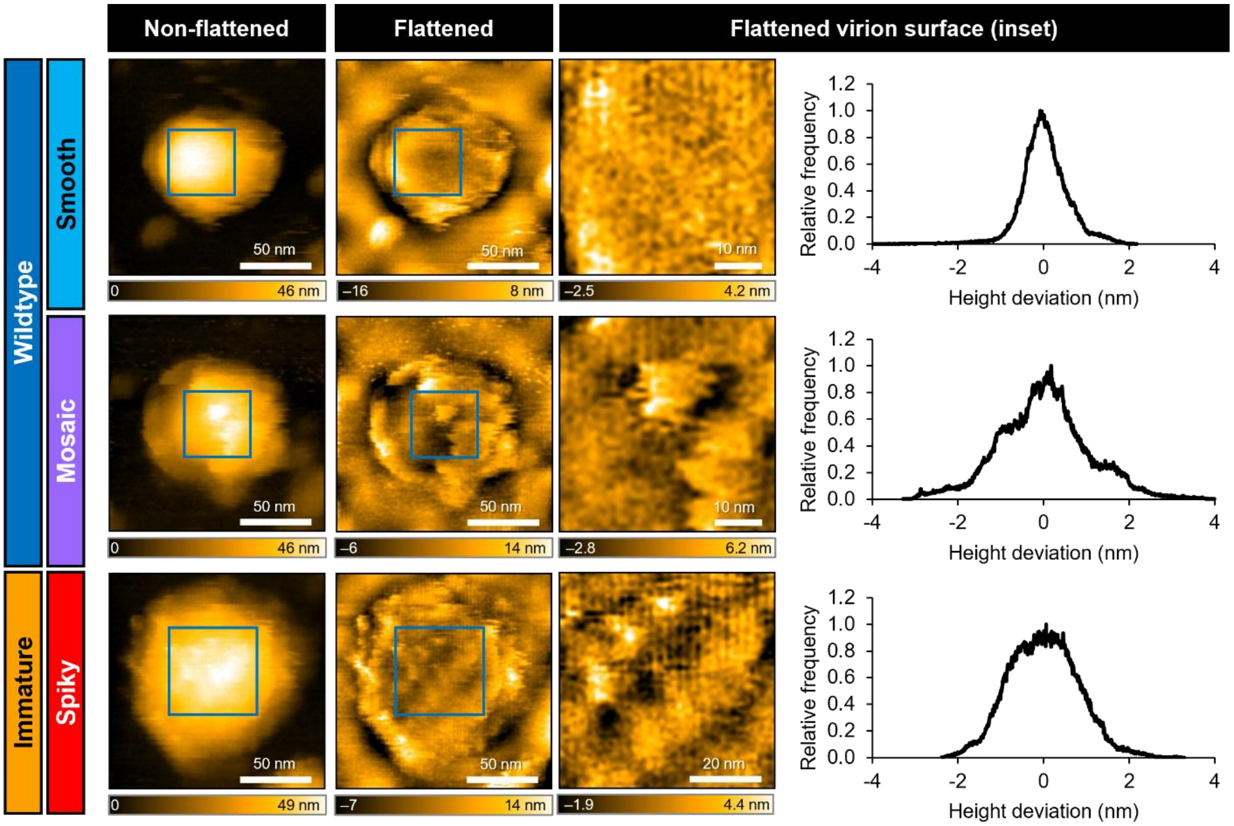


**Supplementary Figure 3. Representative roughness calculation for HS-AFM images.** (Left) Individual HS-AFM images from Fig. 2 are re-shown. (Middle) Images are smoothed with a Gaussian filter ($\sigma$ = 1.0 and then processed by second-order flattening filters (rows and columns respectively) and a first-order flattening filter in the XY plane. (Right) The image is then bounded to the inset box delimiting the upper surface of the virion (45 × 45 $nm^2$, 45 × 45 pixels for MA and PM particles, 60 × 60 $nm^2$, 60 × 60 pixels for IM particles), and the pixel height distribution is plotted. The PM virion shows the highest roughness, indicated by the large height deviation from the flattened plane (SD = 1.05 nm). In contrast, IM is intermediate (SD = 0.80 nm), and MA is the smoothest (SD = 0.58 nm).

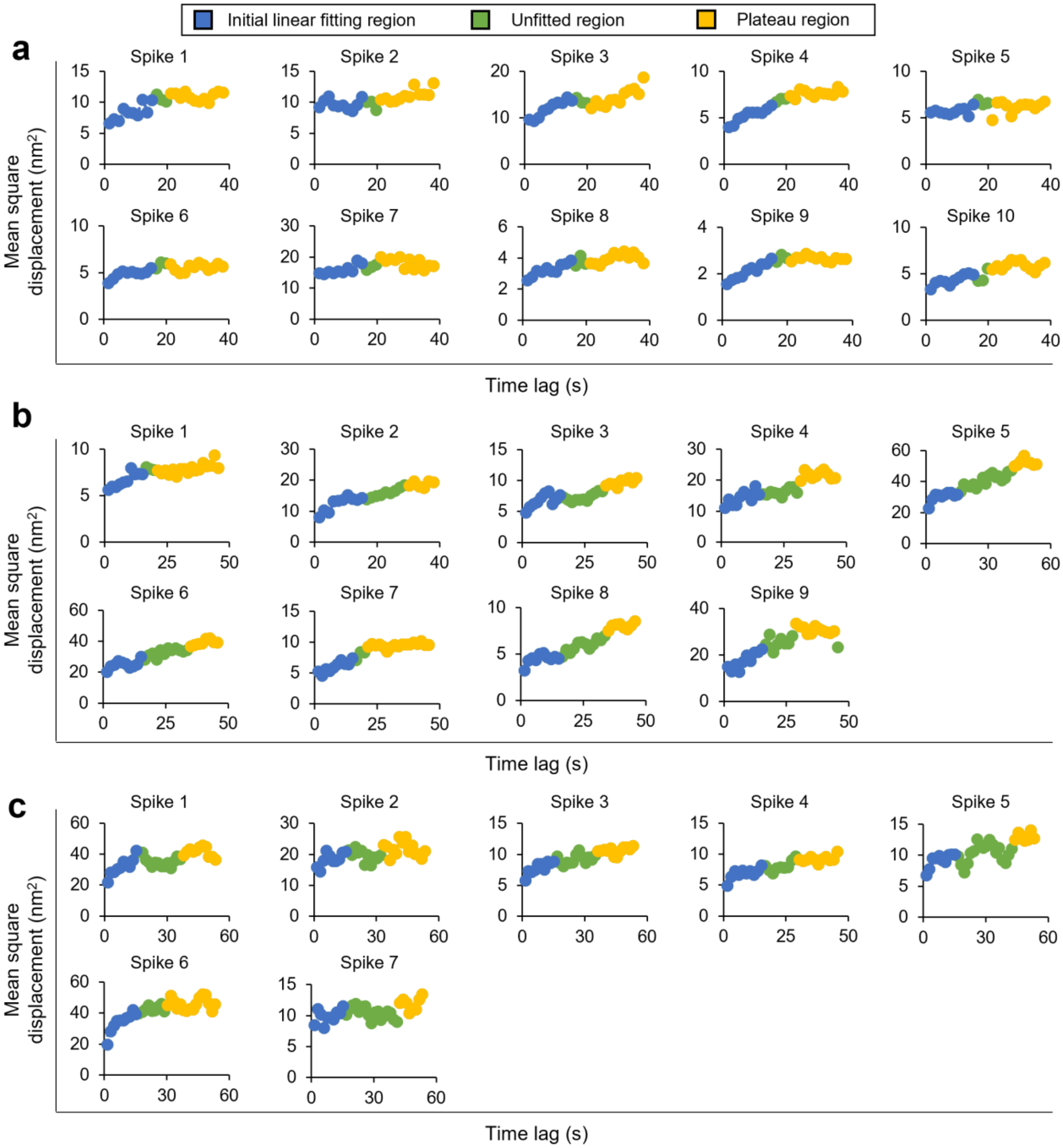


**Supplementary Figure 4. Mean square displacement fitting for determining the radius of confinement of IM spikes.** MSD curves are shown for the particles analyzed in (**a**) Fig. 4, (**b**) Extended Data Fig. 3, and (**c**) Extended Data Fig. 4 respectively. (**a**–**c**) Blue, data used in the initial linear region fitting for determining localization error; green, data not used for fitting; yellow, data used to calculated radius of confinement as described in Supplementary Methods.

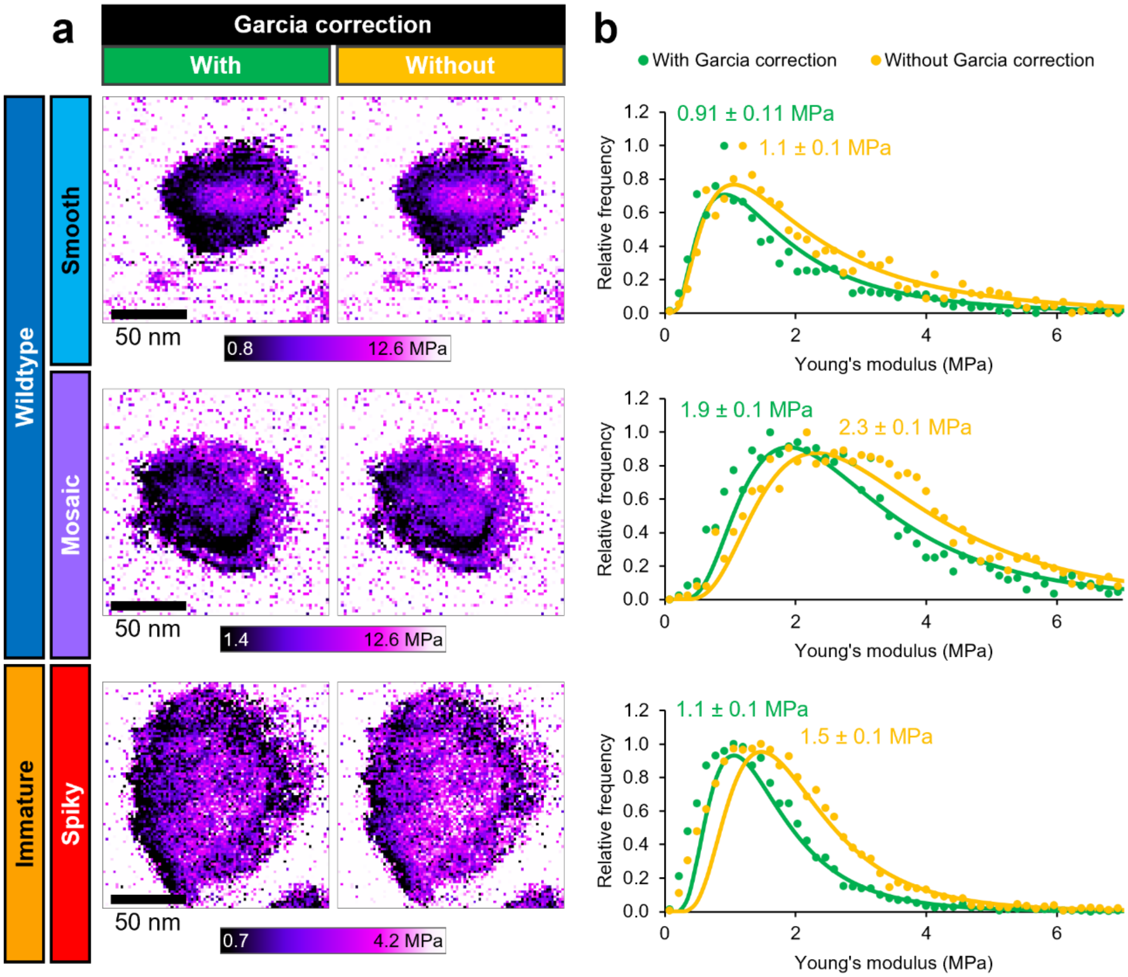


**Supplementary Figure 5. Effect of calculating Young's modulus with the Garcia bottom-effect correction.** (**a**) YM maps for three virions, shown without any filtering. Left and right panels show the map with and without the Garcia bottom-effect correction, respectively, at the same log color scale. (**b**) Lognormal fitting of YM distribution over the virion, with (green) and without (orange) the Garcia correction.

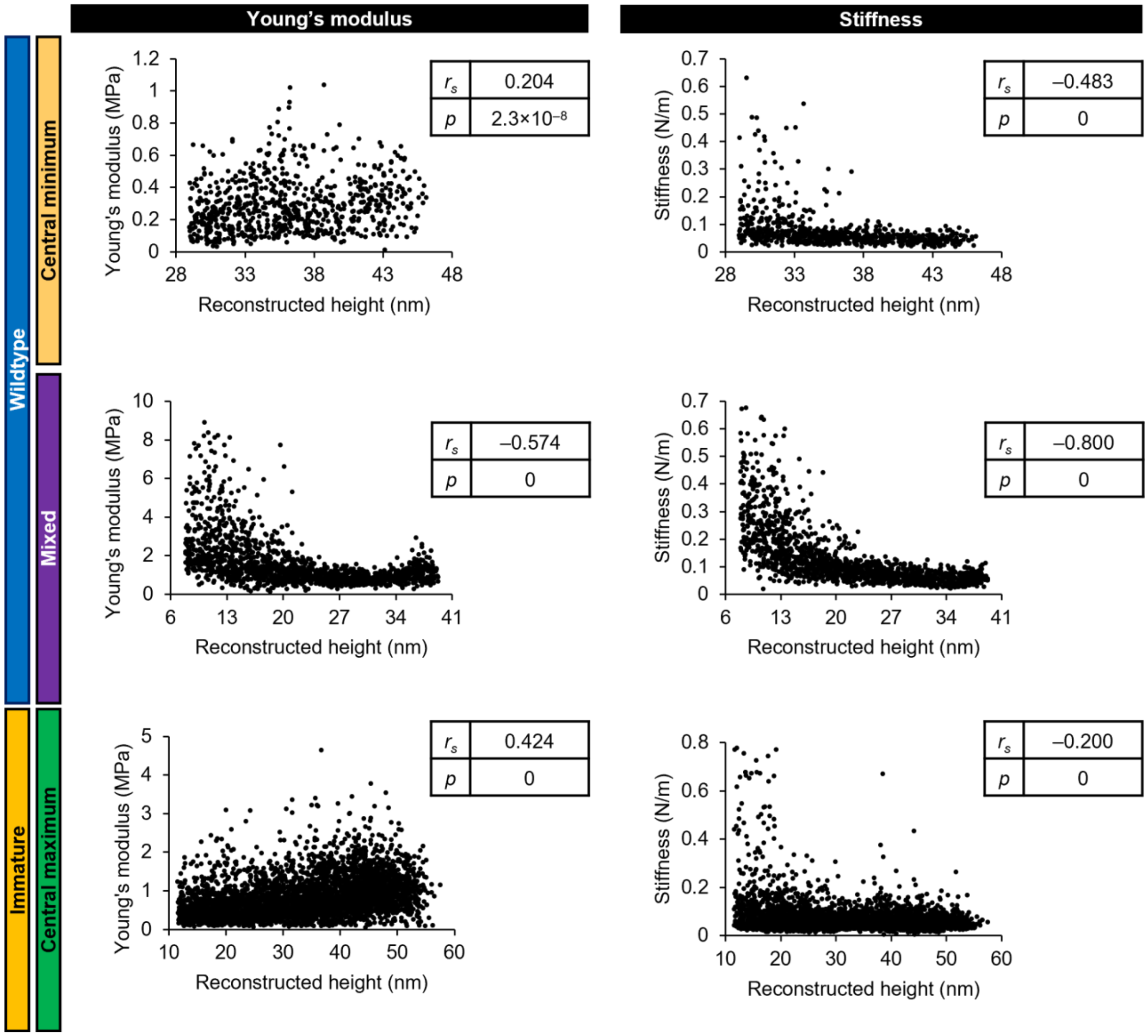


**Supplementary Figure 6. Height dependence of YM and stiffness within single virions.** For the force maps shown in Fig. 5a–c, YM and stiffness are presented as a function of reconstructed height at each pixel (thresholded at 20% maximum height). The Spearman correlation coefficient $r_s$ and corresponding two-tailed $p$ value for each dependency are shown.

### SUPPLEMENTARY VIDEOS

All HS-AFM movies were acquired with the following parameters: scan area 150 × 150 $nm^2$ at 150 × 150 pixels, scan speed 20 µm/s, frame time 1.5 s in only-trace mode. Image processing: tilt corrected, Gaussian filter ($\sigma = 1.0$), median effect filter (1 × 1 pixels). Video processing: stabilized (drift corrected) in ImageJ using StackReg.

**Supplementary Video 1. A representative smooth WT virion.** Z-axis color scaling: black to white, 7 to 47 nm. Video shown at 10 fps.

**Supplementary Video 2. A representative spiky IM virion.** Z-axis color scaling: black to white, 20 to 48 nm. Video shown at 10 fps.

**Supplementary Video 3. A representative mosaic WT virion showing spiky-to-smooth changes.** Z-axis color scaling: black to white, 10 to 48 nm. Video shown at 8 fps. Two scans of the same particle are shown, acquired with an interval of about 5 min between scans.

**Supplementary Video 4. A representative mosaic WT virion showing smearing at the spiky/smooth junction.** Z-axis color scaling: black to white, 2 to 43 nm. Video shown at 10 fps.

**Supplementary Video 5. A representative mosaic WT virion showing relative stability.** Z-axis color scaling: black to white, 4 to 48 nm. Video shown at 7 fps.

**Supplementary Video 6. Real-time tracking of prM–E spike movements on the surface of IM DENV.** Spike maximum traces for spikes #1–10 (as in Fig. 4) are shown with the same colors as in Fig. 4c. HS-AFM was flattened as described in Fig. 4. Z-axis color scaling: black to white, –6.8 to 6.5 nm. Video shown at 10 fps.

**Supplementary Video 7. Topography-mapped nanomechanical parameters reveal maturation-dependent differences.** The topography-mapped projections from Fig. 5 are rotated to enable visualization of Young's modulus, stiffness, and indentation depth across the full virion. Video shown at 30 fps.
